# Prion Protein Folding Mechanism Revealed by Pulling Force Studies

**DOI:** 10.1101/2020.03.09.983510

**Authors:** Theresa Kriegler, Sven Lang, Luigi Notari, Tara Hessa

**Affiliations:** Department of Biochemistry and Biophysics Arrhenius Laboratories of Natural Sciences Stockholm University Svante Arrhenius väg 16C SE-10691 Stockholm, Sweden; Department of Medical Biochemistry and Molecular Biology, Saarland University, Homburg, Germany; Department of Clinical Neuroscience Therapeutic Immune Design Unit CMM, L8:02 Karolinska Institutet, Sweden

**Keywords:** Prion protein, Co-translation folding, Arrest peptide, Xbp1, ER translocation, TRAP complex.

## Abstract

The mammalian prion protein (PrP) engages with the ribosome-Sec61 translocation channel complex to generate different topological variants that are either physiological, or involved in neurodegenerative diseases. Here, we describe cotranslational folding and translocation mechanisms of PrP coupled to a Xbp1-based arrest peptide (AP) as folding sensor, to measure forces acting on PrP nascent chain. Our data reveal two main pulling events followed by a minor third one exerted on the nascent chains during their translocation.

Using those force landscapes, we show that a specific sequence within an intrinsically disordered region, containing a polybasic and glycine-proline rich residues, modulates the second pulling event by interacting with TRAP complex. This work also delineates the sequence of events involved in generation of PrP toxic transmembrane topologies during its synthesis. Our results shed new insight into the folding of such topological complex protein, where marginal pulling by the signal sequence, together with the downstream sequence in the mature domain, primarily drives an overall inefficient translocation resulting in the nascent chain to adopt other topologies.

## Introduction

Protein folding and processing are critical for protein stability and function, and to maintain cellular homeostasis. Protein folding begins when the nascent polypeptide chain is still associated with the ribosome and still undergoing elongation (Fedorov and Baldwin 1997;; Kramer, Ramachandiran, and Hardesty 2001). In eukaryotic cells such “cotranslational” folding occurs for the majority of secretory and membrane proteins, which need to go through the secretory pathway to reach their final destination. The efficiency of this process depends on the conformational stability of the proteins, the mechanisms by which they interact with the translocation apparatus, and on the molecular chaperone repertoire of the specific cell. For a typical efficiently secreted protein, this process initiates with the nascent polypeptide chain binding to the signal recognition particle (SRP), that selectively binds to signal sequences (SSs) and mediates their targeting to the endoplasmic reticulum (ER). Nascent chains bound to the ribosome undergo translocation across the ER membrane via the main protein conducting channel, Sec61complex (herein called translocon), comprised of αβγ subunits (Van Den Berg et al. 2004;; Voorhees et al. 2014). The alignment of the ribosome exit site with the central pore of the translocon is proposed to facilitate direct movement of the elongating polypeptide from the ribosomal exit tunnel across or into the membrane (Becker et al. 2009;; Voorhees and Hegde 2016;; Shanmuganathan et al. 2019). This directs the elongating polypeptide into the ER lumen, cytosol, and lipid bilayer. During this translocation newly synthesized proteins undergo conformational folding, assembly, and post-translational modifications, including signal sequence cleavage, glycosylation and disulfide bond formation (Braakman and Bulleid 2011).

However, not all secretory proteins are equipped with perfectly efficient signal sequences. Depending on the protein to be translocated, i.e. secretory and membrane proteins carrying inefficient SS and less hydrophobic transmembrane domains, the Sec61complex is associated with a range of accessory proteins aiding cotranslational membrane crossing and insertion. Among those, the ‘translocon associated protein’ complex (TRAP), which is comprised of subunits α, β, γ and δ (Wiedmann et al. 1987;; Hartmann et al. 1993;; Pfeffer et al. 2017). TRAP complex has been implicated in facilitating translocation and integration of several model proteins carrying less hydrophobic targeting/insertion domains (Fons, Bogert, and Hegde 2003;; Sommer et al. 2013;; Nguyen et al. 2018). Recent cryo-electron tomography of TRAP–translocon complex has demonstrated the close proximity of TRAP subunits α, β to the Sec61complex and nascent polypeptide chain, while γ and δ subunits have been shown to be bridging between the ribosome and subunits α, β (Pfeffer et al. 2017). Further, proteomics study of TRAP complex dependent translocation of secretory proteins has identified specific features in the signal sequence to be important for recruitment of the TRAP components to the Sec61core complex (Nguyen et al. 2018). In mammalian cells, it has been demonstrated that Sec62 and Sec63 transiently associate with the Sec61 complex (Conti et al. 2015). Sec62/Sec63 has been implicated in the cotranslational ER import of a subset of proteins with less and/or lacking hydrophobic domains (Lang et al. 2012). Sec63 has been shown to be in the vicinity of ribosome-translocon complexes, acting indirectly through Sec62 (Müller et al. 2010;; Wu, Cabanos, and Rapoport 2019;; Jan, Williams, and Weissman 2014), but its role in cotranslational translocation remains to be shown.

Thus, cotranslational translocation is a multistep process that depends on dynamic interactions between the nascent chain and the associated factors. The nature of these interactions suggests secretory and membrane proteins being predisposed to conformational defects, yet structural elements within the precursor proteins responsible for these interactions remain poorly understood. One such example of not perfectly efficient secretory protein, that has shown dependency on ER associated accessory proteins for its efficient translocation, is the mammalian prion protein (PrP). PrP belongs to the glycophosphatidylinositol (GPI) anchor proteins that are targeted to the cell surface, where it is attached to a specific membrane-embedded glycolipid. Structurally, PrP contains an intrinsically disordered N-terminal domain, a helical globular domain in the central portion and a short flexible C-terminal domain that ends with the GPI anchor (Riek et al. 1998). Biogenesis of PrP starts with its import into the ER, where the N-terminal signal sequence is removed, followed by its glycosylation, and its modification by addition of the GPI anchor at its C-terminal end, generating the major secreted form termed ^Sec^PrP. However, PrP has been shown to take two alternative topologies at the ER membrane, termed ^Ctm^PrP and ^Ntm^PrP, where a potential hydrophobic domain (HD) is integrated in the ER membrane. It is established that these two alternatives, together with a significant non-translocated cytosolic PrP (^cy^PrP) population, have direct implications in causing neurodegenerative diseases (Lopez et al. 1990;; Hegde et al. 1998;; Rane et al. 2008;; Hessa et al. 2011).

In this study, to better understand early events of cotranslational folding and translocation of wild type (WT) PrP into the ER, we have determined at which stage during this process the polypeptide chain can start to fold and acquire its native structure or any of the aberrant PrP proteins. In the present study, and based on our previous results, we used the mammalian Xbp1 arrest peptide (AP) as folding sensor, to measure forces generated during cotranslational folding and translocation into ER membrane, and acting on the nascent polypeptide chain (Yanagitani et al. 2009;; Ismail et al. 2012;; Kriegler et al. 2018). We combined this assay with siRNA-mediated gene silencing of translocation components, to identify specific factors, domains, and features governing recognition by translocation apparatus and its accessory proteins. We determined the overall pulling force landscape for PrP folding, and identified at what chain length PrP polypeptides started to misfold into alternative topologies, which can potentially lead to development of diseases. We demonstrated that the mature domains contain features which are important for the translocation process, and which can act in addition to those of the signal sequence. We provide here evidence that a specific hydrophilic sequence in the mature domains of PrP is responsible for recruitment of TRAP complex to the translocon, aiding in its translocation. It is critical to understand the mechanism by which efficient folding is achieved, hence structural elements and specific factors responsible for successful translocation into the ER membrane presents ultimate insight into the molecular basis of prion formation.

## Results

### PrP experiences two pulling events during its translocation

We have recently demonstrated that co-translational translocation efficiency of signal sequences at the ER membrane is primarily dependent on their hydrophobicity, whereas the less hydrophobic sequences, such as the one of PrP, showed inefficient and delayed translocation initiation. In order to understand how folding events are coordinated for PrP, in the present study we addressed the folding and translocation efficiency of full-length wild-type PrP into the ER. We have generated nascent chains trapped at different stages within the ribosome-translocon complex using Xbp1-based arrest peptide (AP). We placed a variant of the Xbp1-AP at a variable distance downstream of each functional domain of PrP known to generate a pulling force on the nascent chain, during cotranslational translocation across ER. These stalled intermediates were representative of the different stages of polypeptide translocation, providing insight into the early stages of folding status (Fig. 1B and Supplementary Fig. 1A and B). PrP possesses a complex domain organization containing a N-terminal cleavable signal sequence followed by a hydrophilic sequence containing a polybasic motif (KKRPK) located directly after the signal peptide cleavage site. Further, it contains an intrinsically disordered N-terminal domain (octapeptide repeat), a weak hydrophobic domain (HD), and C-terminal helical domains, containing two glycosylation sites at positions 181 and 197, that are required for its translocation (Pfeiffer et al. 2013). Finally, a GPI anchor site at position 230 which causes C-terminal cleavage and a S231 linkage to membrane lipids in cells (Fig. 1A) (Miesbauer et al. 2009;; De Fea et al. 1994).

**Figure 1.**
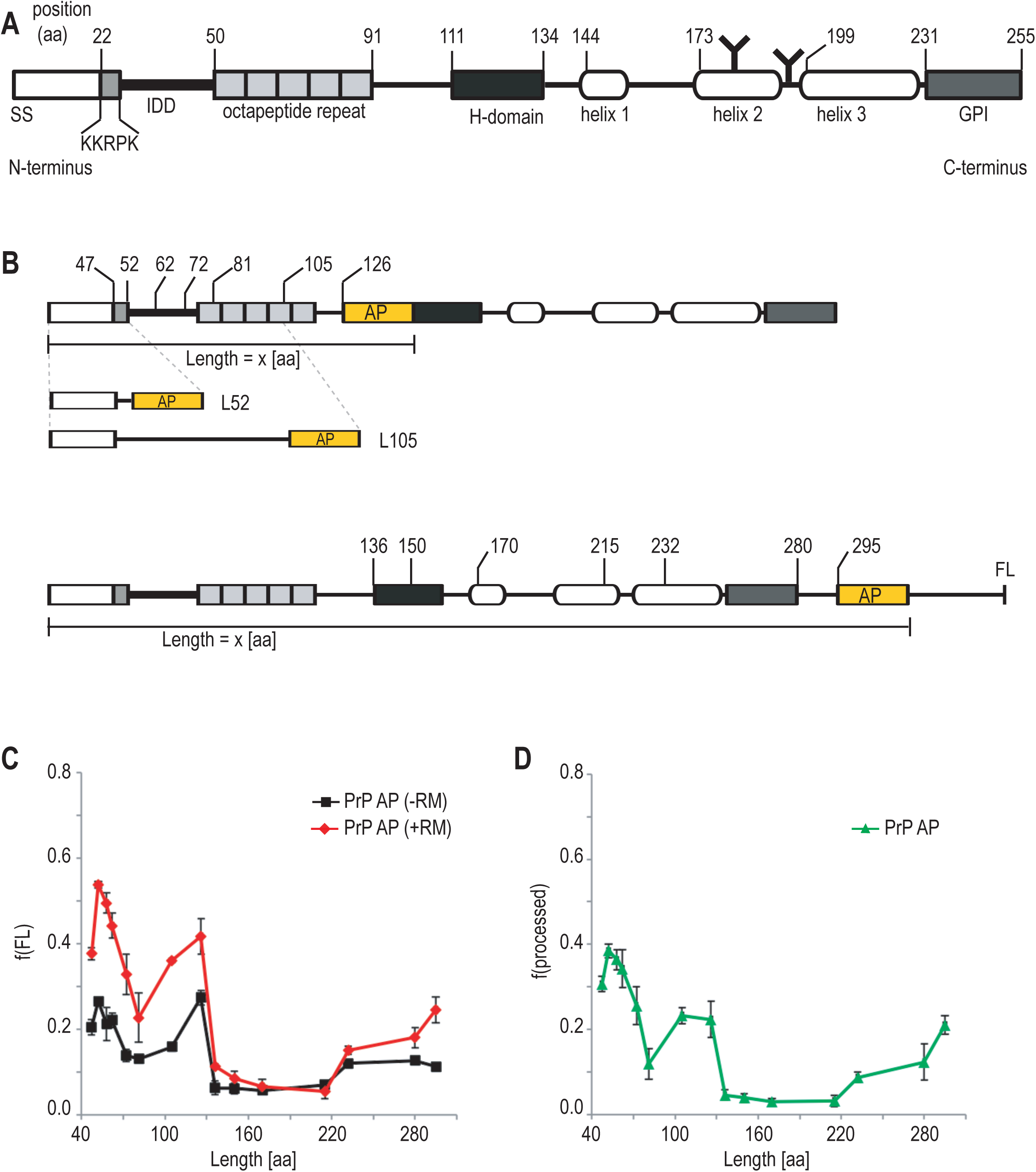
Xbp1 arrest peptide-mediated pulling force profile of Prion protein (PrP) translocation. (A) Schematic overview of PrP features. Signal sequence (SS), Intrinsically disordered domain (IDD), hydrophobic domain (HD) and GPI-anchor sequence and their amino acids (aa) positions are indicated. (B) Representation of the two sets of constructs used in this study. In the first construct set the arrest peptide variant derived from mammalian Xbp1 (AP, yellow) was introduced at position 101 into PrP (see Supplementary figure 1A) resulting in a basic construct with a total arrested length of 126 aa. In the second set of constructs Xbp1 was placed at position 270 including a 15 aa linker after the 255 aa long PrP. An additional extension of 80 amino acids was included after the arrest to create an approximately 9 kDa extension of full length pulled protein for SDS-PAGE separation (see sup figure 1b). This second basic construct has an arrested length of 295 aa. Further constructs were obtained by truncation PCR as indicated by the arrested length and position of AP (see also sup figure 1 a and b). The distance between the start methionine and the C-terminal of the arrest peptide varies in each construct indicated as L (X aa). (C) Pulling-force profile of PrP AP translocation. *In vitro* synthesis of PrP AP in rabbit reticulocyte lysate in presence or absence of dog pancreas rough microsomal membranes (RMs). Fractions of full length (f(FL)) protein (glycosylated, cleaved FL and unprocessed FL) were calculated and plotted in function of L amino acids in the presence (red) or absence (black) of RMs. (D) PrP AP profile showing the fraction of processed FL protein f(processed) plotted in function of L. Fraction f(processed) was calculated as the ratio of the glycosylated + cleaved protein fraction over total protein. The profile shows the mean of at least three experiments, error bars represent their respective standard deviation.

Radiolabeled, fully assembled, translocation intermediates were prepared in rabbit reticulocyte lysate (RRL) based *in-vitro* translation reactions in the presence of [^35^S] methionine and dog pancreas rough microsomes (RM). Upon addition of RMs, targeting and translocation resulted in cleavage of the 22-amino acid SS to produce the mature protein, which was detected at all chain lengths together with glycosylation modifications, which resulted in a band that is resolved at a higher molecular weight than the translation product on SDS-PAGE (Fig. 1B, and Fig. 2, empty arrow head and filled arrow head, respectively). We have previously used several variants of Xbp1-based AP placed at different distances (L) downstream from the start methionine, to allow for better dynamic range (Kriegler et al. 2018;; Shanmuganathan et al. 2019). This range is falling within the criteria of the chain to be pulled from the ribosome PTC site at minimum of 50% and maximum 100% arrest fraction in order to measure a detectable force. For PrP, we used a version of Xbp1 that, in the absence of any pulling force, arrested translation of nascent chains by ∼70% (Fig 1C and 1D). In the absence of RMs each translation product is observed as two bands, representing the unprocessed stalled and unprocessed released PrP;; this later fraction is due to intrinsic leakage (∼30%) in the translation system, where the arrest was overcome, generating full-length protein even in the absence of RM (Supplementary Fig. 1C). Control experiments confirmed that two variants, one containing a stop codon at the arrest site and the other one containing a non-functional arrest peptide, migrated at the corresponding sizes of the arrested and the full-length product respectively (Supplementary Fig. 1C).

**Figure 2.**
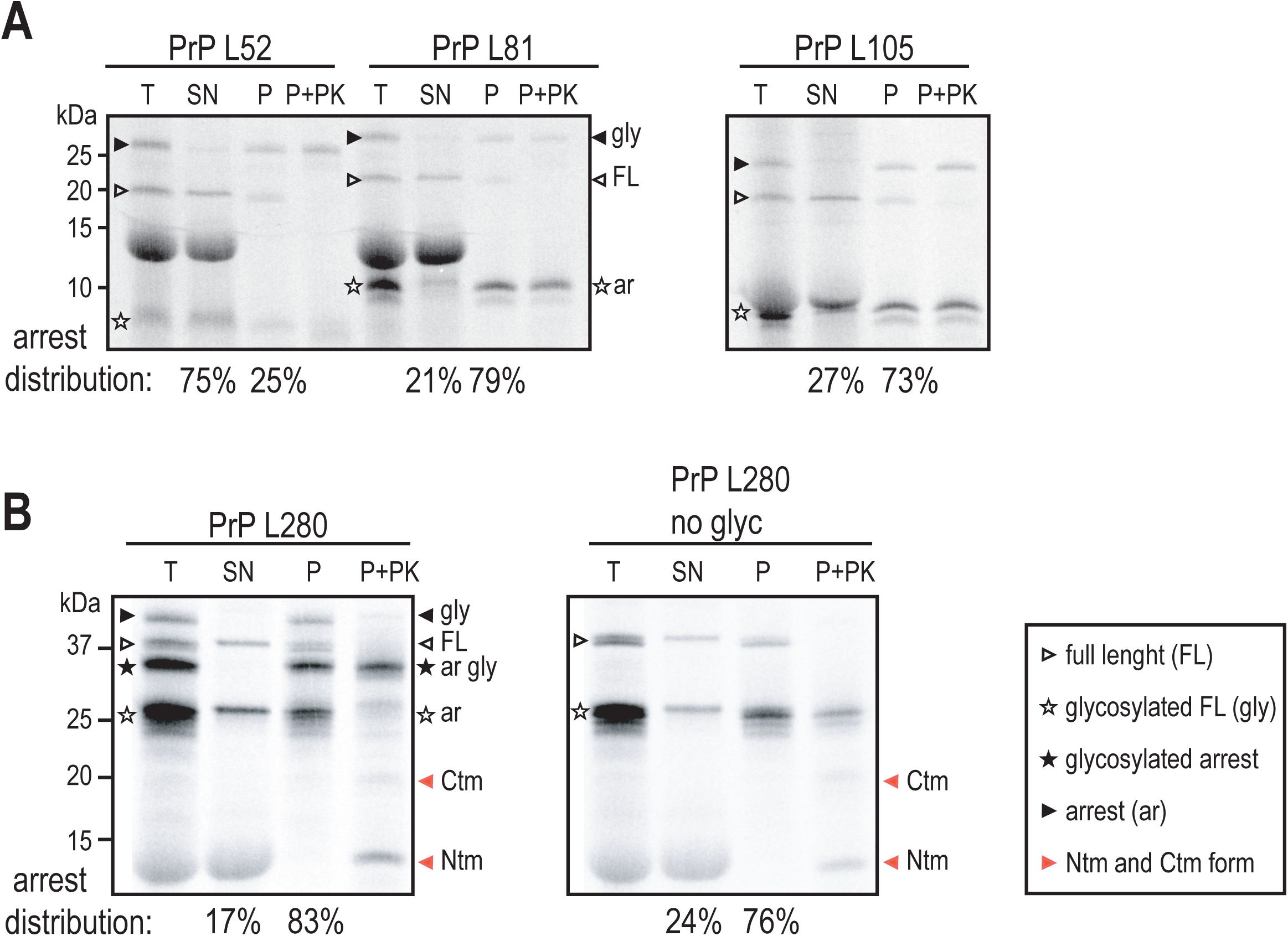
Sedimentation experiments showing increasing membrane association of the arrested population with increased chain length. *In vitro* expressed proteins were subjected to sedimentation on sucrose cushion yielding membrane pellet (P) and supernatant (SN) fractions. The pellet fraction was split into two and either analyzed directly (P) or subjected to proteinase K treatment (P+PK). Full length protein bands are marked with unfilled arrow heads, arrested proteins with empty asterisks and glycosylated protein fractions marked with filled arrow heads or asterisks for full length or arrested populations respectively. Cleaved protein bands are visible for some constructs just below the FL band and not additionally labelled. Percent of membrane bound (P) or un-bound (SN) arrested protein are shown as average from two independent measurements. (A) Sedimentation experiments for short and intermediate chain lengths. (B) Sedimentation experiment for PrP L280 and a mutant without glycosylation sites (PrP L280 no glyc).

Translocation-competent intermediates were assessed from the ratio of the full length released fractions, f(FL), which includes: unprocessed, cleaved, and glycosylated proteins (Full Length, FL), divided by the total synthesized proteins (FL + Arrested, A) (Fig. 1C and Supplementary Fig. 1C). The processed fraction, f(processed), was also calculated as a ratio of cleaved and glycosylated protein relatively to the total translation reaction (Fig. 1D). Pulling force profiles of both f(FL) and f(processed) show that wild-type PrP experiences mainly two membrane dependent pulling events (Fig. 1C and D). The first pulling event was detected at the intermediate length of 52 residues (PrP AP L52), corresponding to about 20 residues being exposed from the ribosome exit channel, and representing an early engagements stage of the SS with the translocon. This is in agreement with our previous results in which PrP signal sequence, fused to prolactin mature domain, was pulled at intermediates lengths L53-L63 and interacted with the translocon components, Sec61α and Sec61β (Kriegler et al. 2018).

The second event was detected as pulling at the intermediate lengths 105 and 126 residues (PrP AP L105, L126), covering exposure of 70-90 residues from the ribosome exit channel, and containing the signal sequence followed by the hydrophilic sequence and the intrinsically disordered octapeptide repeats (Fig. 1C). This indicates a partial folding of this unstructured region and/or the involvement of a folding factor which might contribute to the slight but significant pulling of the nascent chain, as also suggested by the f(processed) profile (Fig. 1D). Interestingly, several studies have demonstrated that irregular structural elements in the mature domain of secretory proteins, elongating within the ribosomal exit tunnel, can modulate the translocation process (Pfeiffer et al. 2013;; Conti et al. 2014). Translocation of PrP was reported to be dependent upon synthesis of helical domains located at residues 172-226, presumably forming the core folding unit (Heske et al. 2004;; Miesbauer et al. 2009). However, only a minor pulling was observed at precursor chain lengths L280 and L295, in which the entire full length PrP nascent chain is exposed outside the ribosome (corresponds to 245 and 260 residues) (Fig. 1C and 1D). This might be due to an additive effect of the helical region folding together with the GPI attachment signal, a marginal hydrophobic segment located at the very C terminus of the protein, being synthesized (Galian et al. 2012).

Next, we confirmed translocation state and processing of PrP by membrane sedimentation experiments to separate membrane bound from the soluble nascent chain intermediates combined with protease K (PK) and glycosylation inhibitor peptide (NYT) treatments. The cleaved glycosylated released FL proteins for lengths L47 through L170 (Fig. 2A, filled arrow heads, and Supplementary Fig. 2A, B) were protected from PK digestion and segregated into the pellet fraction, confirming their translocation in the ER lumen. The uncleaved released FL proteins were instead mostly detected in the soluble fraction (Fig. 2A, empty arrow heads and Supplementary Fig. 2A). We also subjected these samples to glycosylation inhibitor peptide (NYT) treatment to confirm their translocation in the ER lumen (Supplementary Fig. 2A).

The arrested fraction of L52 showed 25 % membrane association that was partially protected from PK digestion, representing early translocation intermediates which are less stably associated with the membrane (Fig. 2A, empty asterisks). The slightly pulled intermediate L105 instead, showed tight membrane associations (70%), protected from PK treatment (Fig. 2A). Further extension of the nascent chains (L280, Fig. 2B and L170 in Supplementary Fig. 2B, see also figure 3B) showed that the arrested population was stably associated (80%) with the membrane fraction and PK protected. Furthermore, we observed signal sequence cleavage of the arrested population starting at chains length 170 to 295, peaking at 215, as demonstrated by PK protection assay. Separation of membrane bound arrested fractions, in the presence of PK treatment, showed the arrest fraction with a cleaved signal sequence to be protected (Supplementary Fig. 2B). This is in agreement with a previous report showing that PrP signal sequence is processed at lengths between 188 and 227 amino acids (Conti et al. 2015).

**Figure 3.**
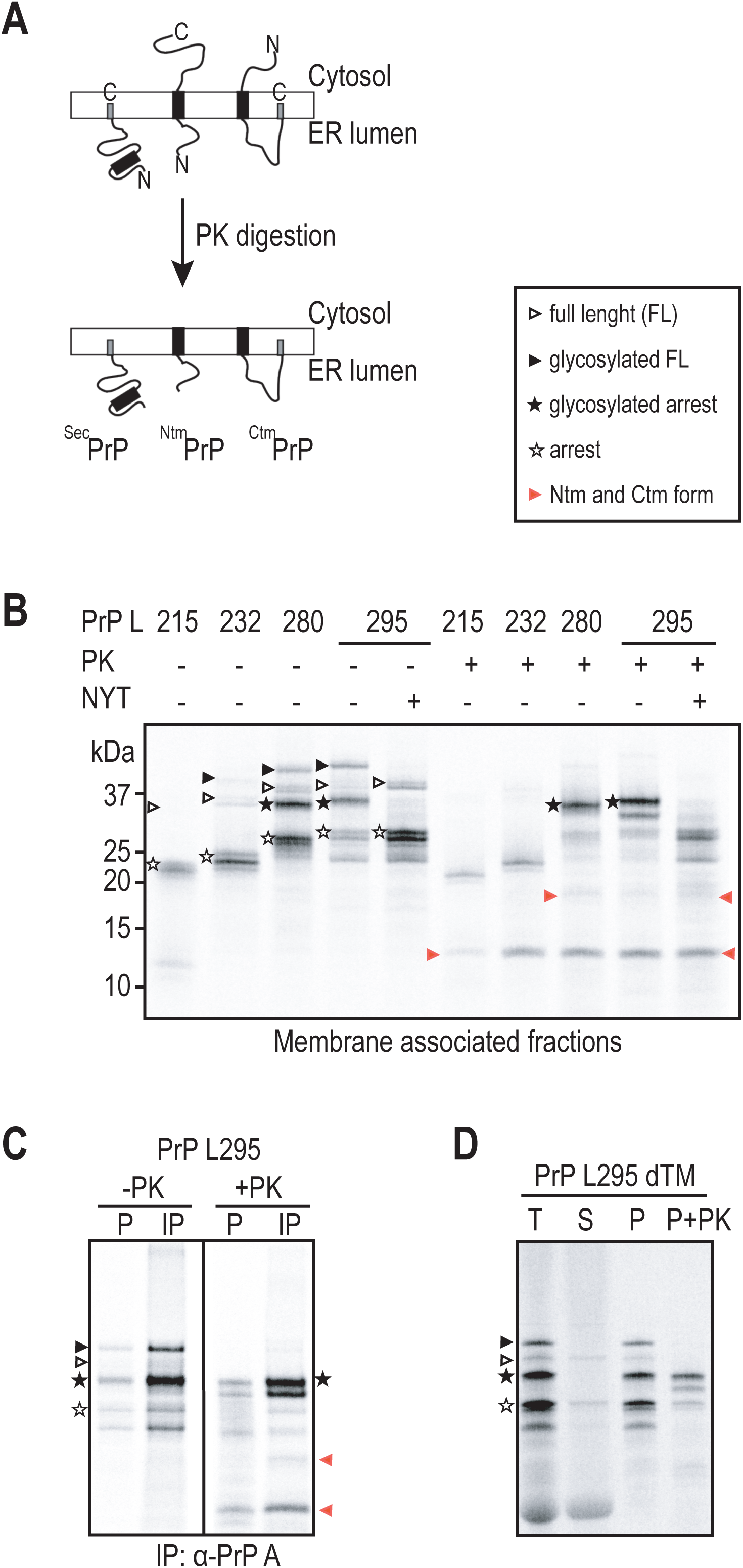
Analysis of PrP AP translocation efficiency across the ER membrane. (A) Schematic of proteinase K assay for topology determination of PrP. (B) *In vitro* expression of long intermediates (L215, L232, L280 and L295, see supplementary figure 2 for L170) in presence of dog pancreas rough microsomes (RMs) was followed by sedimentation on sucrose cushion. Proteinase K treatment (+PK) of the membrane fraction reveal two protected fragments (filled red arrow heads) that corresponds to the ^Ntm^PrP and ^Ctm^PrP forms presented in (A). Unprocessed arrested fractions are indicated with empty asterisks, PK protected glycosylated arrest forms are indicated with a black filled asterisk. (C) Immunoprecipitation (IP) with antibodies (α-PrPA) raised against the N-terminal part of mature PrP (aa 23-38) was performed on PK treated (+PK) and untreated (-PK) pellet samples PrP AP L295 (P). The unprocessed arrested (empty asterisks), Ntm and Ctm forms of PrP (red arrow heads) are indicated. (D) Sedimentation of a PrP variant L295 lacking the potential transmembrane domain (PrP L295 dTM).

Surprisingly, at the chain length L280, the pulled nascent chains, both in the processed (cleaved and glycosylated) and non-processed versions, were membrane associated, but not protected from PK digestion (Fig. 2B, empty and filled arrow head respectively). We found that the arrested population in the pellet fraction was no longer PK protected, but we detected instead three additional PK protected PrP variants, including one with higher and two with lower molecular weights, suggesting that they could have been converted into other known PrP topological forms, the ^Ntm^PrP and ^Ctm^PrP, respectively (Hegde et al. 1998;; S J Kim, Rahbar, and Hegde 2001;; Stewart, Drisaldi, and Harris 2001)(Fig. 2B, filled asterisks and red arrow heads).

To confirm that the higher molecular weight variants were generated by glycosylation of the arrested chains, we used mutant versions of PrP L280 (lacking the glycosylation sites, PrP L280 no glyc). These control experiments clearly showed that the arrested population was represented by polypeptides running at a slightly lower molecular weight, indicating that they were glycosylated and stably associated with the membrane (Fig. 2B, filled asterisks). The other two protected populations appeared instead to run at the same size as the others, suggesting them as not potential substrates for glycosylation (Fig 2B, red arrow heads).

Taken together, our results show that PrP nascent chains experience three pulling events, two main and a third minor event, during translocation across ER membrane: an early pulling event at L52, when nascent chain was in the vicinity of the translocon components, a second event detected at L105-L126,where the nascent chains entered in a more engaged membrane bound state where the unstructured domain was exposed from the ribosome, and a third one, where signal sequence cleavage of the arrested population started at L170 to 215 and glycosylated at intermediate L280. Importantly, we found that the pulled processed FL fraction of L280 nascent chain were not protected in the lumen, despite being modified by glycosylation, and they were converted into the transmembrane forms. These results demonstrate that PrP transmembrane topology formation is dependent upon the timing with which the two topological determinants (the signal sequence and the hydrophobic domain) are presented to the translocation machinery. It has been shown that PrP signal sequence cleavage starts at late stage, at chain length 188 aa, and it was not completed until chain lengths of 227 aa corresponding to the intermediates L215-L295 (Conti et al. 2015).

### Formation of ^Ntm^PrP and ^Ctm^PrP topologies during translocation

Previous studies analyzing its synthesis and translocation have established that PrP can adopt three topological isoforms as part of the normal biosynthesis in the ER. In addition to the main secreted form, ^sec^PrP, with PrP completely translocated across the membrane, two lesser PrP variants, ^Ntm^PrP and ^Ctm^PrP, are inserted as single-spanning membrane proteins with either the N or C terminus translocated into the ER lumen as depicted in Figure 3A (Hegde et al. 1998;; S J Kim, Rahbar, and Hegde 2001;; Stewart, Drisaldi, and Harris 2001). However, from these studies it was not clear when, during polypeptide elongation, these altered topologies started forming.

To address this question, we subjected the longer membrane bound intermediates L215-L295 to PK treatment, and assessed their translocation by following processed arrested fractions (cleaved and glycosylated). Indeed, our results show that, upon digestion with PK, two lower molecular weight bands were protected inside the ER lumen at these intermediate lengths, suggesting that they were partially inserted (Fig. 3B, red arrow heads).

PrP-AP L215 and L232 showed that only the cleaved arrested fraction was protected, since these constructs do not contain any glycosylation sites. At intermediates L280 and L295, the main PK protected fraction was represented by the processed arrested, together with two lower molecular weight bands (Fig. 3B, filled asterisks and red arrow heads). The identity of these bands was confirmed by treatment with glycosylation inhibitor, which resulted in a down shift on SDS-PAGE when compared to the glycosylated arrested population (Fig. 3B). To further confirm the identity of these additional bands as alternative variants of PrP, we performed immunoprecipitation (IP) using specific antibody raised against a region of PrP immediately C-terminal of the signal sequence (residues 23-38) (Emerman et al. 2010;; Ashok and Hegde 2008;; Rane et al. 2008). IP of PrP intermediate L295 revealed that both of the transmembrane PrP forms could be pulled down in the PK treated samples, as well as the glycosylated arrest population (Fig. 3C, red arrow heads and filled asterisks respectively). Further, deletion of the hydrophobic domain (HD) (residues 112–134) from the PrP L295 resulted in abolishment of both lower molecular weight bands confirming them to be the ^Ntm^PrP and ^Ctm^PrP forms (Fig. 3D).

These results indicated the pulled processed population was most likely converted into ^Ntm^PrP and ^Ctm^PrP forms (Fig. 3B, red arrow heads). However, we cannot exclude the possibility that these later forms might be raised from the arrest non-processed fraction as well. This suggest that at these chain lengths PrP were able to get glycosylated, that subsequently deglycosylated before being converted into transmembrane forms. It has been reported that only C-terminal membrane anchorage, by the GPI moiety, affects the efficiency and site occupancy of the N-glycan addition sites of PrP leading to the conclusion that glycosylation and membrane anchorage are co-operative processes (Walmsley, Zeng, and Hooper 2001).

### The second pulling event is depending on mature domain of PrP

Based on our results, the mature domain of PrP experiences a weak pulling at an intermediate length corresponding to the hydrophilic region, following the signal sequence, which contains the highly conserved polybasic sequence (KKRPK) and the intrinsically disordered domain. This prompted us to investigate the role of the mature domains in the translocation of PrP nascent chains. In contrast to signal sequences, few studies have shown that cotranslational translocation efficiency can also depend on mature domains of secretory proteins (S J Kim, Rahbar, and Hegde 2001;; Levine et al. 2005;; Conti et al. 2014).

To investigate the influence of its mature domains, we generated PrP constructs carrying the efficient signal sequence of Prl (Prl-PrP AP), while preserving the mature domains of PrP including the polybasic motif (Fig. 4A). Interestingly, the overall pulling force profile measured for f(FL) and f(processed) showed no change in both pulling events corresponding to the signal sequence and the mature domain, respectively (Fig. 4A). Conversely, placing the same AP version within the coding sequence of wild type full-length prolactin preserved the first pulling peak within the same pulling strength, while completely abolished the second peak, clearly indicating that the mature domains of PrP are responsible for the second pulling event (Supplementary Fig.3A).

**Figure 4.**
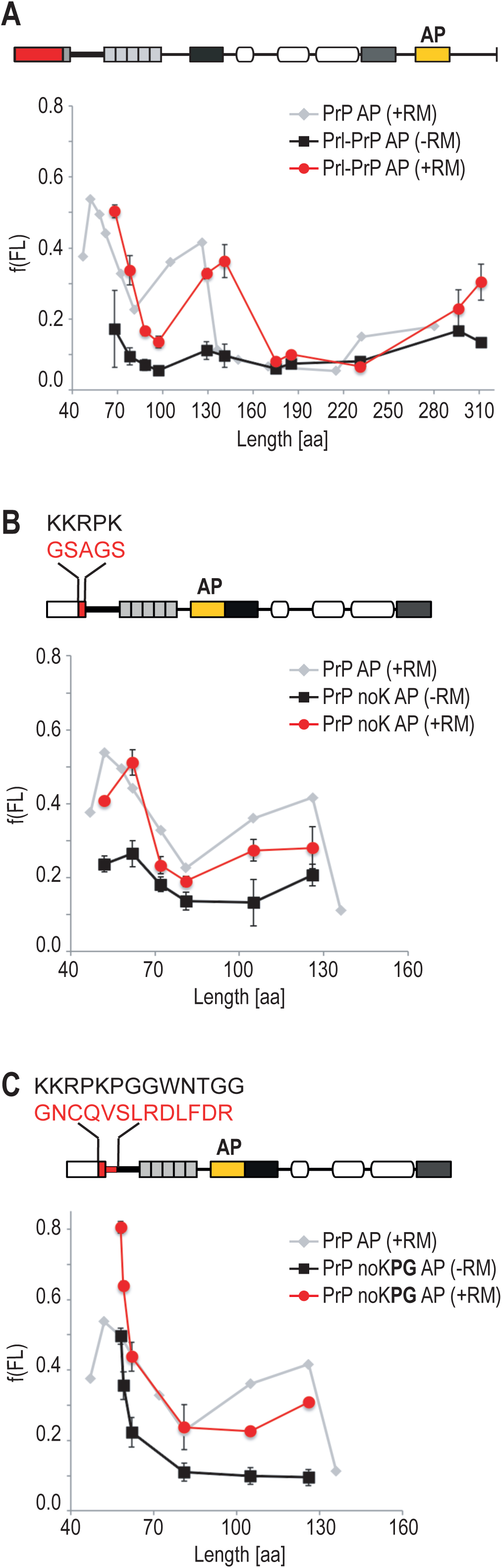
Pulling force experienced during translocation is depended on features of the mature domain of PrP. (A) Schematic representation of the constructs analyzed, top panel. Quantifications of fraction full length (f(FL)) for constructs where the signal sequence (SS) was replaced with the prolactin (Prl) signal sequence (Prl-PrP AP), bottom panel. The calculated f(FL) is plotted as a function of the arrest length of the constructs in presence (red) or absence (black) of dog pancreas rough microsomes (RMs). For comparison, the profile obtained for PrP AP in presence of RMs is plotted in light grey. The means of at least three experiments are shown (except for L175, L185 and L231, where n=2), error bars represent their respective standard deviations. (B) Pulling-force profile for PrP mutant with replacement of 4 lysines (noK) with an uncharged sequence as indicated. The f(FL) data for PrP noK AP constructs is plotted in function of the arrested construct length in presence (red) or absence (black) of RMs. The profile shows the mean of at least three experiments, error bars represent their respective standard deviations. For profile of processed protein only see sup figure 3. (C) The first 12 aa of the mature domain of PrP (starting with aa 23) that are enriched in positively charged amino acids as well as glycine and proline, were replaced with the first 12 aa from mature Prl (PrP noK**PG** AP). The graph shows the f(FL) in presence (red) and absence (black) of RMs over the arrest lengths. Data point are plotted as average of at least three experiments, error bars represent their respective standard deviations. For profile of processed protein see Supplementary figure 3.

Because charged residues in close proximity to the signal sequence cleavage site may affect translocation (Fujita et al. 2011) we next examined the role of the hydrophilic sequence directly following the signal sequence in PrP translocation. To this end, we generated two series of mutations in the constructs with chain lengths L52-L126. In the first series, we replaced the poly basic residues (KKRPK) with an uncharged sequence (PrP noK AP) and in the second one an additional GP-rich stretch (KKRPKPGGWNTGG) was replaced with unrelated sequence from Prl sequence (PrP noK**PG** AP). Both of these substitutions resulted in a slight decrease in the f(FL) of the second pulling peak for both L105 and L126 intermediate lengths (Fig. 4B and 4C), while the f(processed) remained unchanged (Supplementary Fig. 3B and 3C). Interestingly, we noticed a significant increase in the first pulling peak for a construct with the combined substitutions, even though the non-membrane dependent pulling (no RM) also increased. These results indicated a sequence specific pulling, dependent on charges and hydrophilic residues, playing an important role in efficient translocation of PrP mature domains. Moreover, abolishment of these hydrophilic residues seems to have weakened interactions between PrP and a presumably assisting factor(s).

### Semi-permeabilized cells allow silencing components of the translocon complex that affect pulling events

To gain insight into the mechanism and factors responsible for exerting the pulling force on the PrP nascent chain, we sought to reconstitute the pulling force measurement in human cells. We analyzed translocation of stalled PrP intermediates using semi-permeabilized cells (SPC) pretreated with targeted siRNA to deplete specifically various translocation components (Lang et al. 2012). In short, HeLa cells were incubated with the siRNAs or the non-targeting control siRNA and then treated with digitonin for permeabilization. Equivalent amounts of cells were used for the pulling force assay. Radiolabeled PrP translocation intermediates were synthesized in RRL and the reactions were supplemented with the different semi-permeabilized siRNA-treated cells as a source of ER membrane. Stalled translation intermediates were assessed for signal sequence cleavage and glycosylation modification to determine their translocation and folding status.

In initial experiments, we established translocation of PrP nascent chain and measured pulling force of the PrP AP intermediates used in the RRL based in vitro experiments. Fractions full-length f(FL), and processed f(processed), were calculated in the same way as for the in vitro experiments (see material and method). Overall the cell-based system recapitulated the translocation efficiency faithfully, and the force profile showed similar pulling events experienced by the PrP nascent chains, with lower sensitivity, supporting the conclusions from the in vitro studies using RRL (Fig. 5A and Supplementary Fig. 4A and 4B).

**Figure 5.**
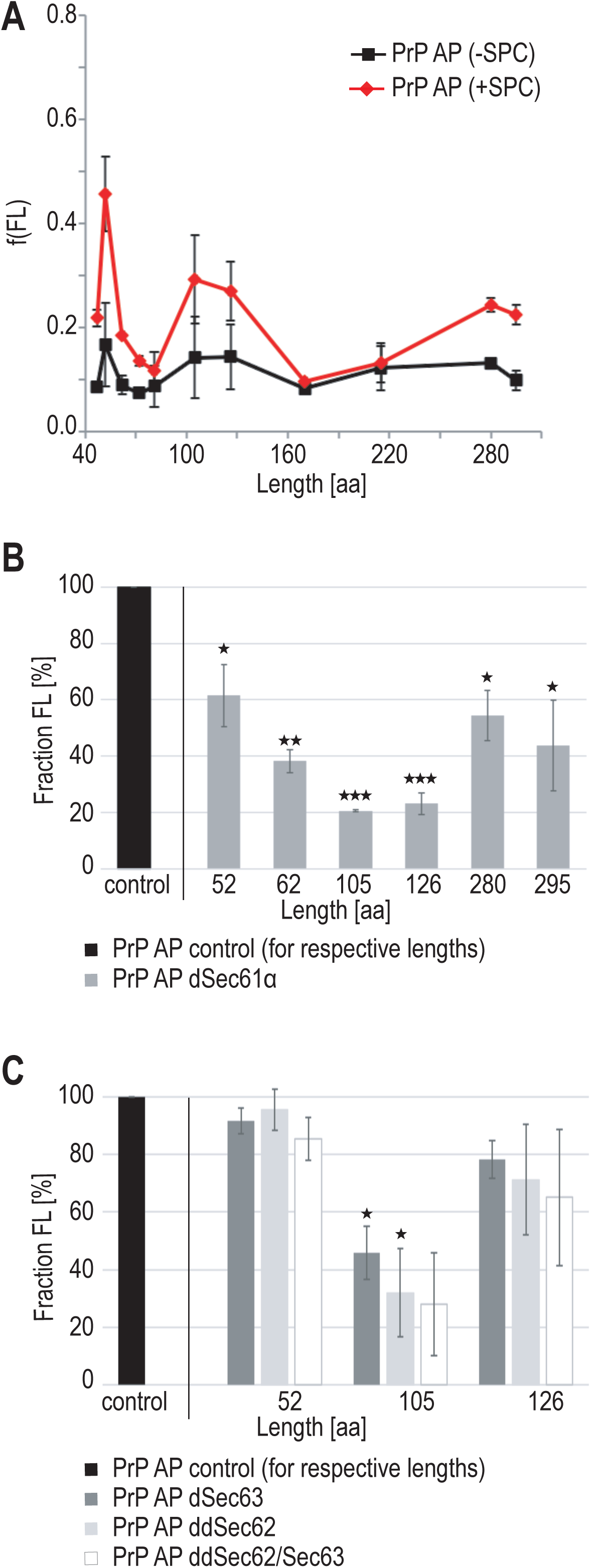

Previous study has provided clear biochemical evidence that PrP nascent chains were in a stabile complex with the several translocon accessory components such as TRAP complex and Sec62/Sec63 (Conti et al. 2015;; Pfeffer et al. 2017). We next adapted the SPC system to determine which factors were affecting the pulling of the PrP nascent chains by depleting the translocation components, Sec61α and associated accessory proteins, Sec62/Sec63 and Trapβ.

In general, the resulting membranes from digitonin permeabilized Hela cells were substantially depleted (by 75% - 90%) relative to control siRNA treated membrane (Supplementary Fig. 4C). As a proof of concept, we started by assessing the silencing of the *SEC61A1* gene (Sec61α) using a Prl AP construct series from our previous published study, where Prl AP chain length L74 was pulled maximally in a Sec61α dependent manner (Kriegler et al. 2018). Pulling of these control constructs, Prl AP L57-L80, was decreased (by 74%) in *SEC61A1* silenced SPC attesting the validity of experimental approach in capturing changes in pulling profile upon silencing of specific component (Supplementary Fig. 4D). This also supported our previous conclusion that Prl was translocated efficiently in a Sec61complex dependent manner (Kriegler et al. 2018).

Pulling of PrP AP intermediate L52 and L62 were modestly reduced (by 40% and 60%, respectively) representing the yet non-fully engaged and Sec61 dependent intermediates, translocation of intermediate lengths L105 and L126 were reduced by 80%, representing the maximum reduction in pulling capacity by this siRNA set (Fig. 5B). At precursor lengths L280 and L295 depletion of Sec61α resulted in a reduction by 60%, reflecting the fact that pulling was also exerted on these longer intermediates, and probably reflecting a slightly less dependence on Sec61α for their translocation. Interestingly, this decrease was largely preserved upon replacing the PrP signal sequence with that from Prl (Prl-PrP AP) (data not shown). Overall these experiments highlight the strict requirement for the translocon core component, Sec61α, for efficient PrP translocation across the ER membrane independently of the signal sequence.

Next, we assessed whether Sec62 and Sec63 depletion affected PrP pulling force profiles at any intermediate lengths. Hela cells were treated with siRNAs directed against the non-coding (SEC62-UTR) regions of the SEC62 gene and the non-coding (SEC63-UTR) regions of the SEC63 gene or the negative control siRNA (Greiner et al. 2011;; Lang et al. 2012). Western blot analysis confirmed the almost complete silencing (by over 90%) after 96 hours treatment (Supplementary Fig. 4C).

Translocation of PrP intermediates into Sec63 depleted SPC membranes were mostly affected for the second pulling peak (mostly L105), which was decreased by 60% and 50% (Fig. 5C), while the effect on the first pulling peak was minor. Similarly, depletion of Sec62 gene individually or in combination with Sec63 gene affected mostly the second pulling event (Fig. 5C). These results illustrates that Sec62 and Sec63 proteins contribute to a moderate pulling of PrP nascent chains L105 and L126, which correspond to the second pulling event in the PrP pulling profile.

### Silencing of TRAPβ reduces the second pulling event in a signal sequence-dependent manner

TRAP complex (α, β, γ and δ) has been implicated in assisting cotranslational translocation of PrP as well as a subset of secretory and membrane proteins in a signal sequence dependent manner (Fons, Bogert, and Hegde 2003;; Sommer et al. 2013;; Nguyen et al. 2018). It was indeed shown that it can interact with nascent chains at late translocation stages (Wiedmann et al. 1989) and has been demonstrated to physically associate with Sec61 (Pfeffer et al. 2017).

We examined the role of TRAP complex in translocation of PrP nascent chains by depletion of TRAPβ subunit. We used a previously established TRAPβ-targeting siRNA that resulted in an efficient reduction and degradation of the other subunits of TRAP-complex without affecting the Sec61α protein levels (Pfeffer et al. 2014;; Nguyen et al. 2018). Silencing efficiency of TRAPβ subunit was confirmed by western blot (Supplementary Fig. 4C).

While the pulling of intermediates L52 was not affected by TRAPβ knock-down, we observed a reduction in the pulling of the intermediate L62 by 45% (Figure 6A). Strikingly, silencing of TRAPβ resulted in 80% reduction of the f(FL) of both intermediate lengths of the second pulling event. We also observed that the pulling for the longer precursor lengths (L280 and L295) were reduced almost by half indicating that the minor pulling events observed for PrP at these lengths were also depending on Trap for their translocation.

Although, protein depletion via targeted siRNAs was not 100% optimal, we could nonetheless detect a clear diminishing effect on the second pulling event and a modest effect on the PrP translocation at chain lengths L62, L280 and L295 indicating the timeline for TRAP complex dependent pulling starting as early as L62, a maximum at L105-L126 and a continue pulling until translation of the nascent chain is completed.

This data is in agreement with previous experiments in which TRAP complex could be cross-linked with nascent chains at late translocation stages (Wiedmann et al. 1989). To analyze whether this diminishing effect was depending on the signal sequence and/or the mature domain of PrP, we used the Prl-PrP AP mutant construct carrying a more efficient signal sequence. Prl-PrP AP showed similar reduction levels of the second pulling event (L129 and L144) providing evidence that TRAP complex was crucial for pulling PrP mature domain during its translocation (Fig. 6B).

**Figure 6.**
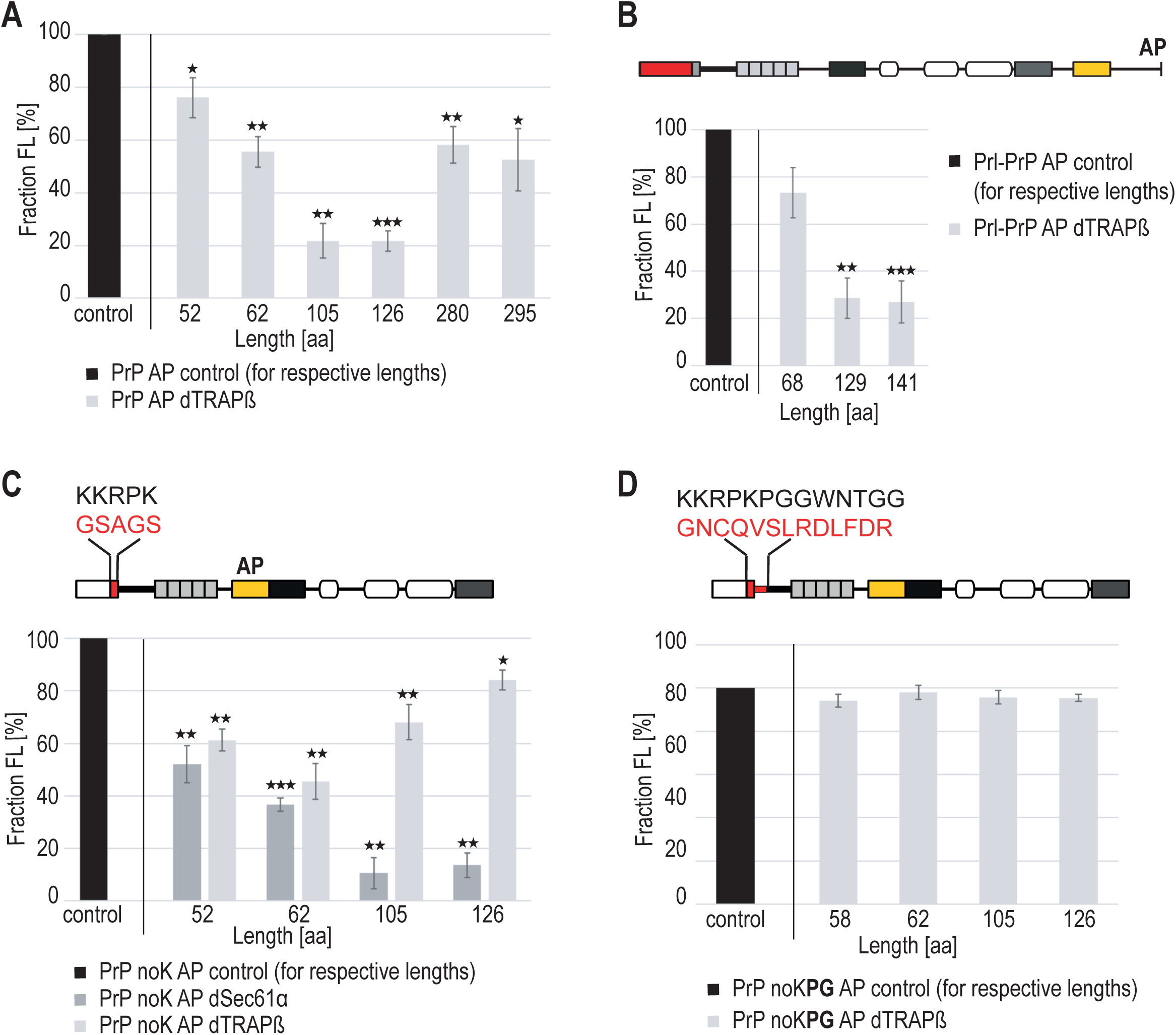

To characterize features in the PrP mature domain associated to the abolishment of the second pulling peak, we used the two mutation series of the chain lengths L52-L126 from figure 3. Upon TRAPβ depletion, the f(FL) for intermediates lengths L105 and L126 of PrP noK AP was almost restored to its normal levels, increased to 65% and 87% respectively, abolishing the Trap dependent pulling effect (Fig. 6C). In contrary, we noticed modest decrease of the f(FL) levels, more TRAPβ dependent, for intermediates lengths L52 and L62 suggesting the polybasic residues might play a role in delaying PrP signal sequence translocation at those early chain lengths (Fig. 6C). However, when we tested the second mutant series, PrP noKPG AP, that had the GP-rich residues replaced, the levels of the f(FL) were restored completely to the control siRNA levels at all intermediate lengths (Fig. 6C). Further, depletion of Sec61α was not affected per se by these mutations, indicating that decrease in pulling force was TRAP specific effect (Fig. 6C).

Taken together our data provide evidence for sequence specific TRAP-depended pulling acting on PrP nascent chains. This sequence specific pulling affected (decreased) primarily the second pulling event, while the first pulling event seemed to be at least partially more dependent on TRAP and Sec61α in the absence of the charged cluster. This also indicated that the signal sequence per se seems to have minor influence on the second pulling event consistent with the notion that TRAP complex intervene at this stage in a sequence specific manner.

## Discussion

A crucial step in the biogenesis of most secreted proteins is their cotranslational translocation across ER membrane. During this step the N-terminal signal sequence interacts with the translocon allowing it to open and insert the mature domain of the protein through the translocon into the ER lumen (Voorhees and Hegde 2016). Efficient signal sequences such as the one of Prl ensure translocation of the elongating nascent chain through the translocon before sufficient polypeptide emerges into the cytosol to stably fold (Walter and Blobel 1981). In the case of PrP, this process seems far from optimal due to its complex topology. PrP carries a signal sequence that is not efficient in initiating translocation, and the putative HD domain promotes formation of neurotoxic topological variants, ^Ctm^PrP and ^Ntm^PrP, or, in the case of failed translocation, it leads to the generation of misfolded cytosolic PrP (Hegde et al. 1998;; Soo Jung Kim and Hegde 2002;; Miesbauer et al. 2009;; Hessa et al. 2011).

In this study, we addressed the folding and translocation efficiency of PrP into ER membrane using Xbp1-AP mediated ribosomal stalling (Ismail et al. 2012;; Kriegler et al. 2018). Xbp1-AP functions as a sensor to detect cotranslational nascent protein folding via the translocon. The interaction between the translocon and the nascent chain will generate a force that represents not only PrP nascent chain folding, but also the binding of partners to the nascent chain outside the ribosome, in the translocon environment. Folding and translocation status are here evaluated by limited proteolysis (PK digestion) and glycosylation modification assays.

We show that PrP nascent chains undergo two main pulling events, at chain intermediate lengths 52 and 105-126 amino acids, and a minor one when the full-length polypeptide translation is completed. The first pulling event is representing an early engagement stage with the Sec61 complex, where the signal is cleaved and the arrested ribosome nascent chains appear to be partially membrane-bound and partially protected by the ER bilayer (Kriegler et al. 2018). The second event represents pulling of the hydrophilic sequence and the intrinsically disordered domain, as judged by signal cleavage and translocation into the ER lumen. Previous studies have found that a polybasic region within the hydrophilic sequence is critical for protection against PrP aggregation and consequent neurodegeneration (Turnbaugh et al. 2011). The intrinsically disordered domain contains octapeptide repeat, and it has has been proposed to bind copper or zinc ions on the cell surface, governing cellular ion content (Pauly and Harris 1998). Similar studies have shown that zinc induced mature domain folding occurs within the ribosome exit tunnel/translocon junction (Conti et al. 2015;; Nilsson et al. 2015). This folding will then prevent further translocation of the mature domain to ensure unfolded/partially folded nascent chain passage into the lumen, thus in agreement with the pulling event observed when such a domain is exposed from the ribosome. In our domain swap experiments in which signal sequences are exchanged between PrP and Prl, we here clearly demonstrate that the second pulling event is not dependent on the signal sequence, but rather on the mature domain itself. These findings demonstrate an interplay between a functional signal sequence and structural properties of its downstream mature domain supporting the proposed model by Conti and colleagues, where the nascent chain does not pass immediately from the ribosome through the translocon after ER targeting, but rather accumulates transiently within the ribosome/translocon junction (Conti et al. 2015).

Surprisingly, we detect no pulling exerted on the nascent chains exposing the marginally hydrophobic domain (HD, L170) nor the C-terminal helical domain located at residues 172– 226 (L215 and L232), but rather a weak pulling is detected once translation of the full-length is completed at L280 (Fig 1B and C). However, we cannot exclude the possibility of this weak pulling might be also due to the marginally hydrophobic GPI signal (Galian et al. 2012). Our results indicate that a sufficient force is sensed by the N-terminal domain to pull more than 80% of the total nascent chains across the membrane. Thus, we can conclude that PrP translocation is mainly dependent on its N-terminal domain, contradicting previous study showing that translocation of the intrinsically disordered N-terminal domain is depending upon synthesis of C-terminal helical domains (Pfeiffer et al. 2013).

Further, signal sequence cleavage of the arrested population starts at chain lengths L170, corresponding to after the emergence of the HD domain from the ribosome, and is not completed until chain length L280 when glycosylation happens. This suggests that PrP nascent chains begin to translocate at a chain length of 135 aa, and become fully committed at a length of 200 aa, corresponding to helical domain emergence from the ribosomal exit tunnel. Interestingly, we found that the pulled processed fraction of L280 lost its glycosylation and was not anymore protected in the lumen;; nascent chains were instead converted into the transmembrane ^Ctm^PrP and ^Ntm^PrP forms. Deletion of HD in PrP abolished the transmembrane formation and delayed the signal sequence cleavage (Conti et al. 2015).

More important, these results identify the time point, intermediate length L 215 to L295, during which PrP nascent chains deviates from its normal biogenesis and starts forming the neurotoxic transmembrane topologies.

The pulling force profile was here recapitulated in mammalian cells by using semi permeabilized Hela cells as source of ER membrane, suggesting that similar interactions and factors are conserved between the RRL in vitro and cell based systems. Next, we set out to determine which factor (s) is responsible for the observed pulling on the PrP nascent chains;; we depleted the translocation components Sec61α and the associated accessory proteins, Sec62/Sec63 and TRAP complex by gene silencing approach (Lang et al. 2012). The binding of such factors may play roles in providing a chaperone-like role particularly for proteins carrying the less efficient signal sequence and/or intrinsically disordered proteins during membrane translocation(Lang et al. 2012;; Conti et al. 2015). Depletion of the core translocon subunit Sec61α markedly decreases the second pulling peaks intermediates, and modestly decreased the first one, for the intermediate L52, by 50%, reflecting the not yet fully engaged state of nascent chain with the translocon (Voorhees and Hegde 2016;; Kriegler et al. 2018). A number of studies implicated PrP nascent chain to relay on Sec62/Sec63 complex and BiP chaperone for its efficient translocation (Haigh and Johnson 2002;; Ziska et al. 2019;; Young et al. 2001). PrP nascent chains, lacking either the HD or GPI anchor, expressed in Sec63 knock-out cells failed to be translocated across the ER membrane compare to the wild type cells (Lang et al. 2012;; Wilson et al. 1995). Further, PrP nascent chain longer than 160 aa was found to stabilize a complex containing Sec61/62/63 at the ER membrane (Conti et al. 2015).

However, depletion of either Sec62 and Sec63, or combined, did not results in any major effect on the pulling force profile as opposed to the depletion of the other factors. This suggests that Sec62 and Sec63 may merely have a supportive role in stabilizing translocon complex during cotranslocation of polypeptides containing weak determinants, but do not have active role in pulling the nascent chain. Another explanation could be due to the fact that these factors seem to interact transiently with PrP nascent chains during late stage in their synthesis and the method used here is unable to capture such weak pulling events.

On the other hand, we found a striking effect on the pulling force profile upon depletion of TRAPβ. Our data suggested that the second pulling events exerted on the PrP mature domain was due to interaction with the TRAP complex. This sequence specific TRAP-depended pulling effect seemed to be at least partially due to the charged cluster and GP-rich stretch that immediately follow PrP signal sequence. While the signal sequence per se seems to have minor influence on the second pulling event or in this context. Recent proteomic study has identified features in the signal sequence of a subset of secretory protein containing GP-rich sequence to be dependent on TRAP complex for their efficient translocation (Nguyen et al. 2018). Our results provide new insights into the important role of TRAP complex in assisting PrP translocation beyond the signal sequence and again relying on the presence of a G/P rich motif.

## Acknowledgements

We like to thank Dr. Ramanujan Hegde for the generous gift of the prion protein antibody. This work was supported by grant from the Swedish Foundation for Strategic Research to TH.

## Author contributions

TH conceived the project, TH and SL designed the study. TK and SL performed the experiments and evaluated the data together with TH. LN, TK and TH wrote the manuscript. All authors commented on the manuscript.

## Material and Methods

### Plasmids and Constructs

All Prion protein (PrP) constructs were based on the hamster PrP sequence in SP64 vector that has been previously described (Hegde et al. 1998). A variant of the 25 amino acids long arrest peptide from the X-box binding protein (Xbp1) was introduced into the PrP sequence at two different positions. For all short constructs the starting construct contained the arrest peptide (AP) at position 101 replacing a stretch of 10 amino acids (KPS-site) as shown in Figure 1B and supplement figure 1A. For longer constructs the AP was introduced with a 15 amino acid linker at the end of the protein sequence and an additional extension to distinguish the arrested from the full length protein form on SDS-PAGE (Figure 1B and supplement Figure 1B). The Xbp1 arrest peptide, a less functional arrest peptide, or a stop codon (Kriegler et al. 2018) were introduced in the constructs by Gibson assembly (Gibson et al. 2009), the additional linker was also introduced using Gibson assembly, while parts of the C-terminal extension were created by extended oligos in overlap PCR. All the oligomers used to create the basic constructs are shown in Table 1. Shorter constructs counting from the start to the end of the AP were obtained by PCR using forward and reverse primers that were complementary to the regions indicated in supplemental figure 1A with right-facing and left-facing arrows, respectively. Oligo sequences are presented in Table 2. Overlapping PCR was the method of choice, but in some cases, restriction-ligation was required, in which case the phosphorylation of the oligos was performed by using the T4 Polynucleotide Kinase from Thermo Fisher, according to manufacturer’s protocol, followed by ligation using T4 ligase from the same manufacturer.

**Table 1.**
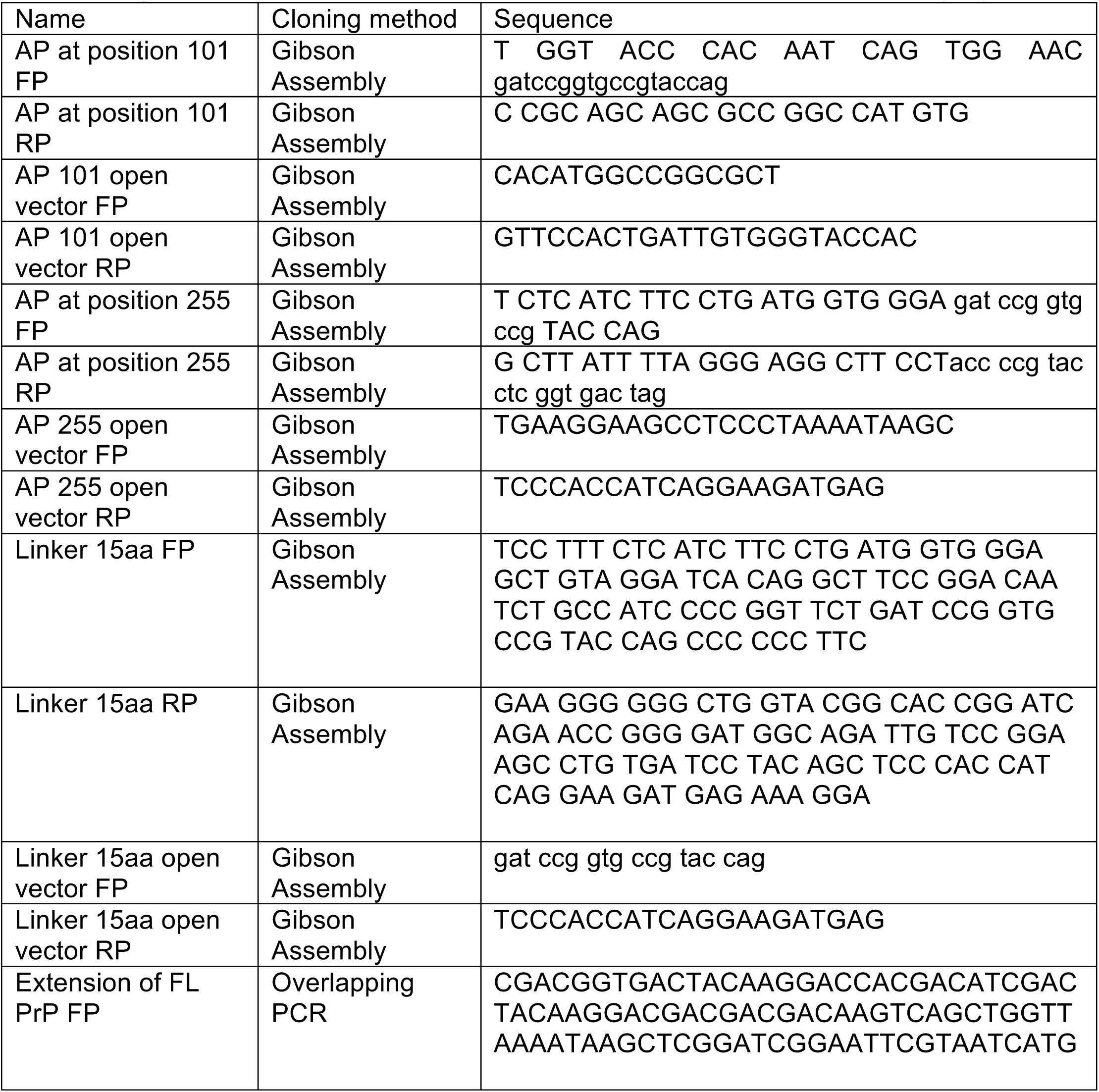
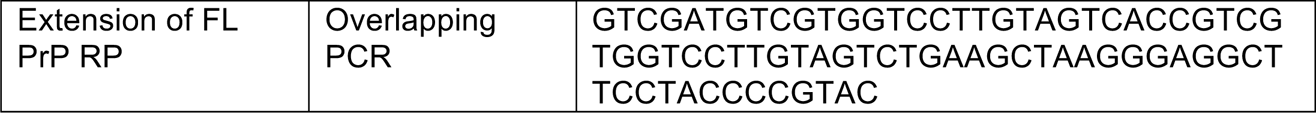
Oligos to create basic PrP constructs with Xbp1 derived arrest peptide (AP)

**Table 2.**
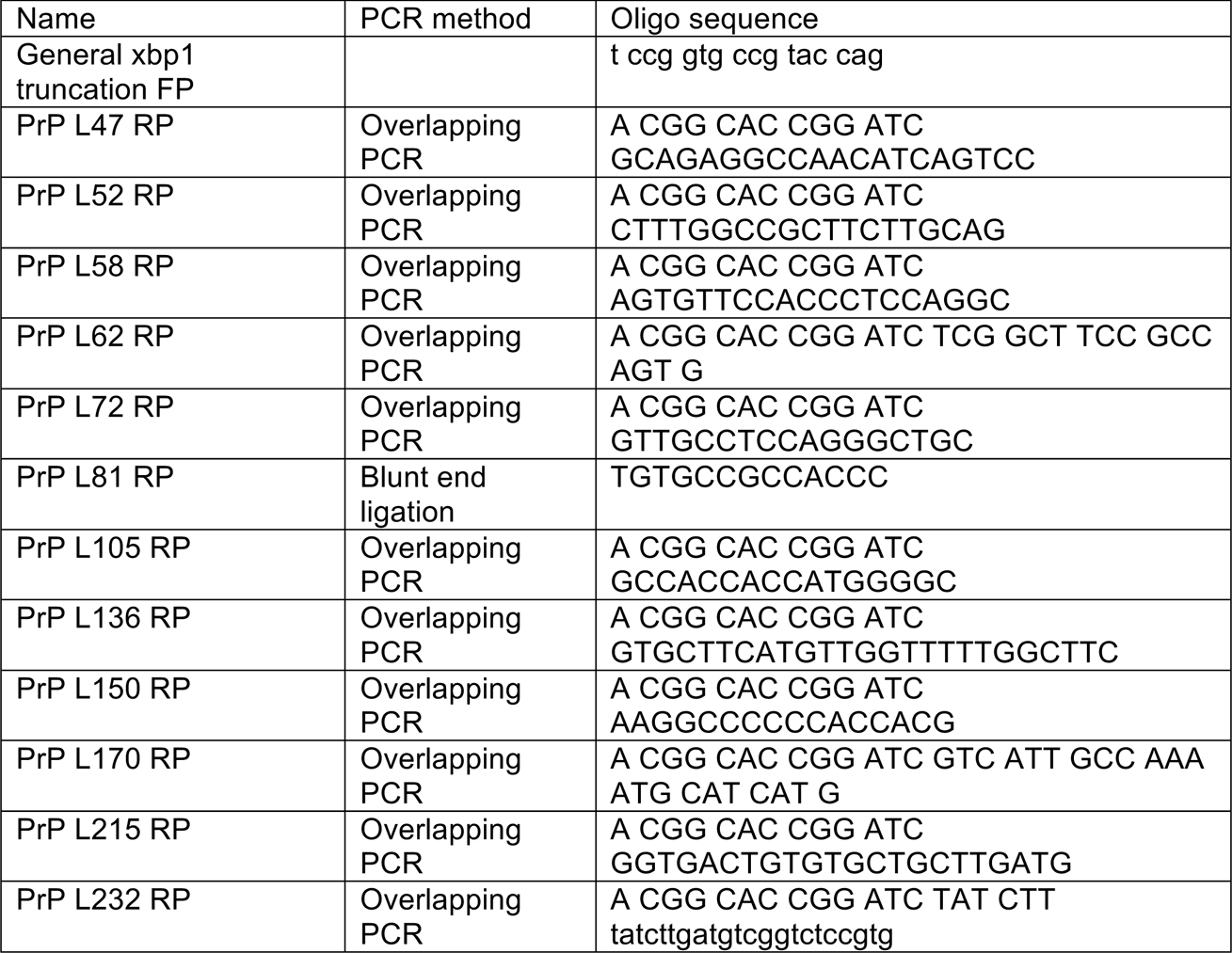
Oligomers to create truncated lengths or PrP AP constructs

Control constructs with PrP SS in Prl background and Prl containing Xbp1 have been described previously (Kriegler et al. 2018). Constructs with Prl SS in PrP background have been generated using overlapping PCR using the oligomers in Table 3. All further modifications have been introduced by site directed mutagenesis or overlapping extended PCR using oligomers shown in Table 3. All constructs were transformed into chemically competent DH5-α cells.

**Table 3.**
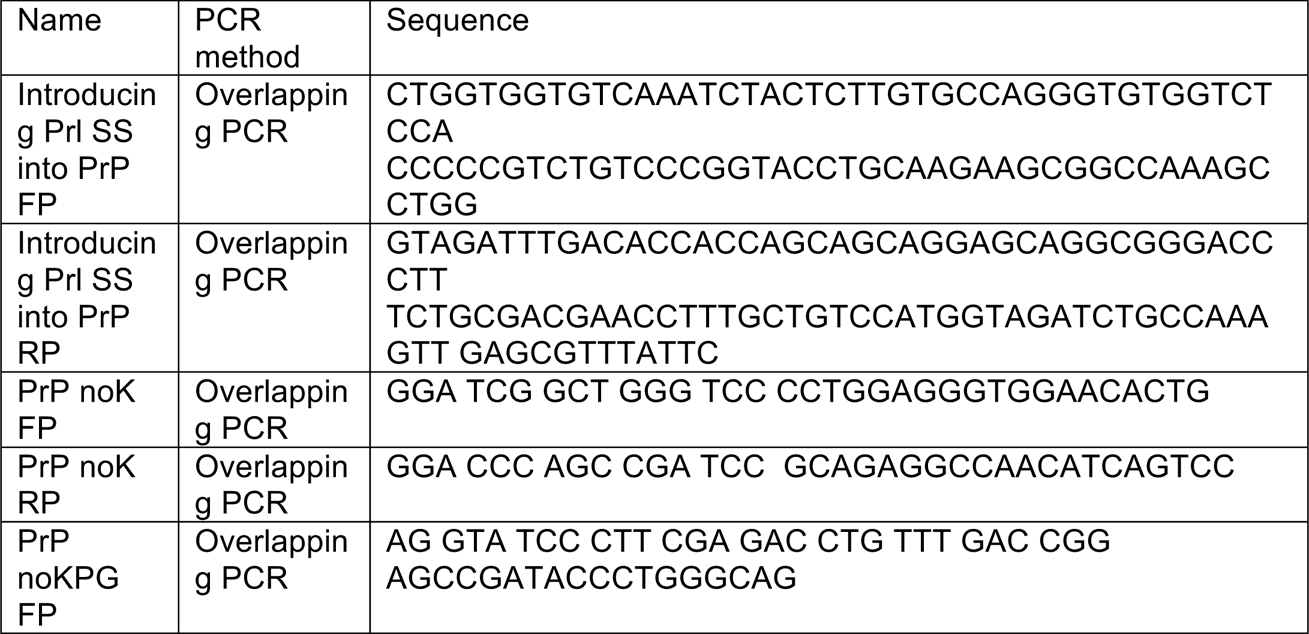
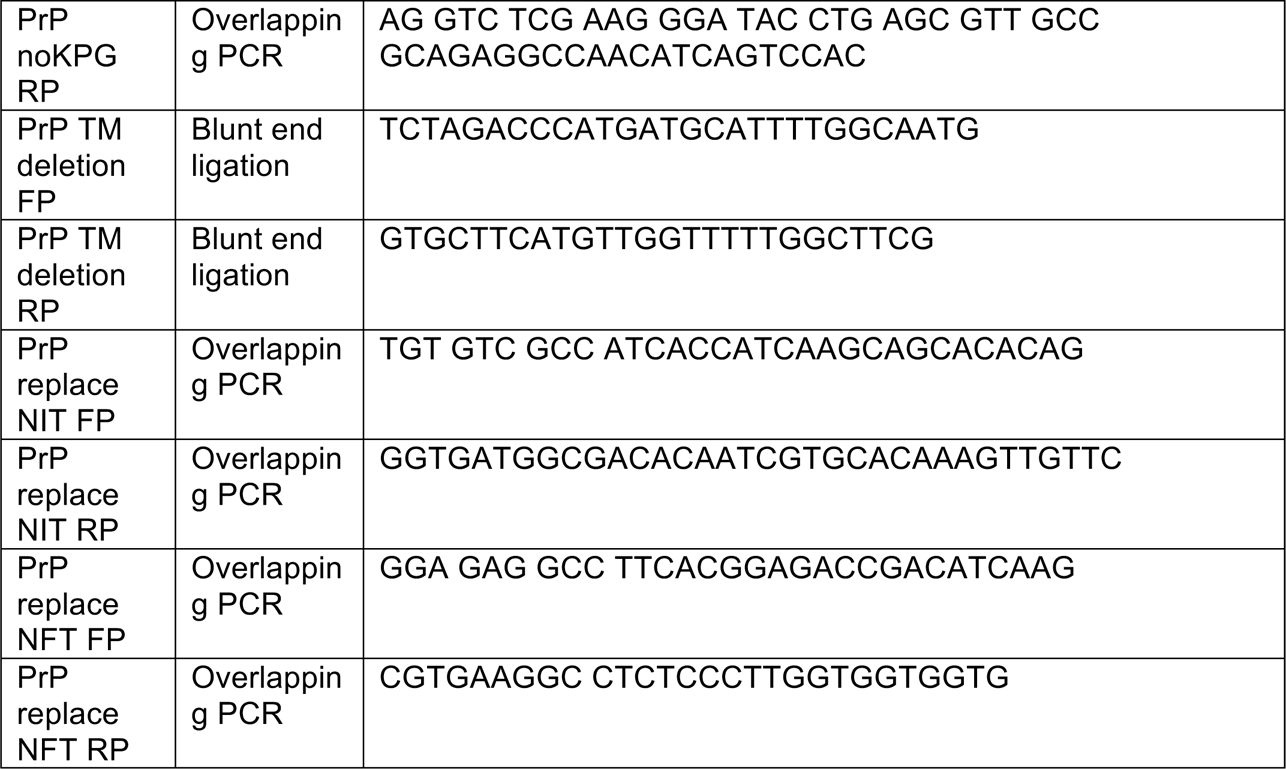
Oligomers for mutated variants of PrP AP

### Cell culture

Short HeLa cells (ATCC no. CCL-2) or regular HeLa cells (DSM no. ACC57 from the German Collection of Microorganisms and Cell Cultures) were cultivated at 37 °C and an atmosphere containing 5% CO_2_ in Dulbecco’s modified Eagle’s medium (DMEM, Gibco) containing 10% fetal bovine serum (Sigma), 4mM L-glutamine (Sigma), 1mM sodium pyruvate (Sigma) and 1% penicillin/streptomycin (Sigma).

### Preparation of siRNA treated semi-permeabilized cells (SPCs)

As described before (Ziska et al 2019) 5.6 x 10^5^ cells were seeded in 4 ml DMEM in a 6-cm culture plate and transfected with targeting or control siRNA (Table 4) (from Qiagen) to a final concentration of 20 nM for each siRNA used, using HiPerFect transfection reagent (Qiagen). After 24 h of normal culture incubation, medium was replaced and cells were transfected a second time. For all siRNA used for this study total incubation time for the depletion was about 96 hours. Silencing efficiency was confirmed by western blotting using rabbit antibodies raised against human Sec61α (1:2500 dilution), Trapβ (1:500 dilution), Sec62 (1:500) and Sec63 (1:500 dilution), that were kindly provided by Dr. Sven Lang. The antibodies were visualized using goat anti-rabbit peroxidase conjugated IgG (1:10000) (Thermo Fisher scientific (Cat # 31402)) and the Azure biosystems c600 imaging system (with cSeries Capture Software, version 1.9.8.0403) and analyzed with Fiji open source image processing software (version 2.0.0-rc-65/1.51u). Preparation of digitonin SPCs was according to published protocols (Ziska et al. 2019;; Haßdenteufel et al. 2018;; Lang et al. 2012) with final cell numbers of 40 000 cells/μl.

**Table 4.**
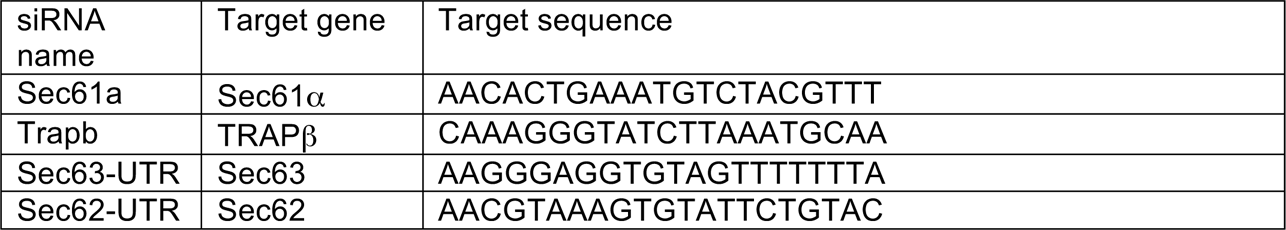
Sequences of siRNAs used in this study.

### In vitro transcription/translation

All basic transcription-translation reactions for PrP constructs were performed using a Rabbit Reticulocyte Lysate (RRL) derived SP6 coupled system (Promega) for 30 min at 30 °C in the presence of 2.5 μCi [^35^S]-methionine in a total volume of 10 μl. Reactions were performed with addition of 0.5 μl column washed microsomes (RM) from dog pancreas (tRNA products) when indicated. The reactions were stopped by addition of sample buffer and boiling for 5 min at 95 °C before treatment with RNaseA (15 min, 37 °C). Reactions containing SPCs were performed in a total of 5 μl containing 1.5 μl SPC suspension (40 000 cell/ μl) or Potassium-Hepes-Magnesium buffer (KHM) for controls. They were incubated 30 min at 30 °C as for the basic reactions. The SPC reactions were then chilled on ice and centrifuged for 20 min at 4°C and 15000 rpm in a benchtop centrifuge. Pellets were resuspended in sample buffer and boiled for 5 min at 95 °C. Samples were separated by SDS-PAGE on 12% or 10% Tris-glycine polyacrylamide gels and analyzed by autoradiography.

To inhibit N-glycosylation the transcription/translation reactions were performed in the presence of the tripeptide NYT (0.1 mM final concentration).

Due to low expression levels, the Prl and Prl-PrP control experiments in the SPC system were performed in an uncoupled RRL derived system (Sharma et al. 2010) starting from purified PCR products. Transcription was performed with an SP6 polymerase (New England Bioscience) for 1 h at 40 °C. Transcripts were then added to the translation reaction containing 1.25 μCi [^35^S]-methionine in 5 μl reactions and 1.5 μl SPCs when indicated. Sample preparation, RNase treatment, visualization and analysis were as described above.

### Sedimentation on sucrose cushion

The selected constructs were incubated in RRL for 30 min at 30 °C. Samples (20 μl total volume) were loaded on a 80 μl sucrose cushion (low salt: 250 mM sucrose, 100 mM KAc, 50 mM Hepes pH 7.9, 5 mM MgAc;; high salt: 250 mM sucrose, 500 mM KAc, 50 mM Hepes pH 7.9, 5 mM MgAc) and were centrifuged for 4 min at 46 000 rpm at 4°C in a Optima MAX-XP Ultracentrifuge with a TLA 100.3 rotor (Beckman). The pellet was suspended in (20 µl) sucrose suspension buffer (50 mM Hepes 7.9, 250 mM sucrose, 100 mM KAc, 5 mM MgAc) and analyzed directly or treated with proteinase K. For digestion reactions, pellets were incubated with proteinase K (0.5 μg/μl final concentration) for 1 hour on ice (Görlich and Rapoport 1993), then the reactions were stopped by adding phenylmethylsulfonyl fluoride (17 mM final concentration). All samples were resolved by SDS-PAGE and analyzed as described above.

### Immunoprecipitation

Where indicated, sedimentation pellet samples were subjected to immunoprecipitation (IP) before or after PK treatment. The selected samples were mixed with an equal volume of 2% SDS solution (in 0.1 M Tris pH 8) and boiled for 5 min at 95 °C. After chilling at room temperature, the samples were diluted with cold IP-buffer (50 mM Hepes pH 7.4, 300 mM sodium chloride and 1% Triton X-100) to a final volume of 1 ml. They were then incubated with PrP A antibody overnight at 4°C, with end-over-end rotation. Equilibrated protein A-beads (Bio-Rad) were added to each sample and incubated for 2 hours rotating at 4°C. After extensive washing of the beads, the samples were eluted in 1x sample buffer (62.5 mM Tris, pH 6.8;; 2% SDS;; 10% glycerol;; 100 mM DTT and 0,002% bromphenol blue) for 5 min at 95°C.

### Quantification and statistical analysis

Autoradiography images were acquired using a FLA-9000 phosphorimager equipped with ImageReader software version 1.0 (Fujifilm Corporation). Band intensities were quantified using Fiji (ImageJ open source software, version 2.0.0-rc-65/1.51u) and analyzed using EasyQuant (in-house developed quantification software based on QtiPlot). Fractions of full length (FL) protein was calculated as previously shown by using:

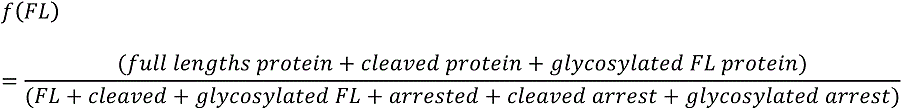

All data was confirmed in at least three independent experiments if not indicated differently. All line graphs show the average of the different experiments with the standard deviation as error bars (if not indicated differently). All bar plots show the average of all values relative to the respective control experiment (set to 100%).

For statistical relevance a one-sided paired t-test was performed on all data obtained from the silencing experiments using student’s distribution values to evaluate p-values. P-value levels are always stated in the figure legends when indicated.

## Legends for Supplementary figures

**Supplementary figure 1.**
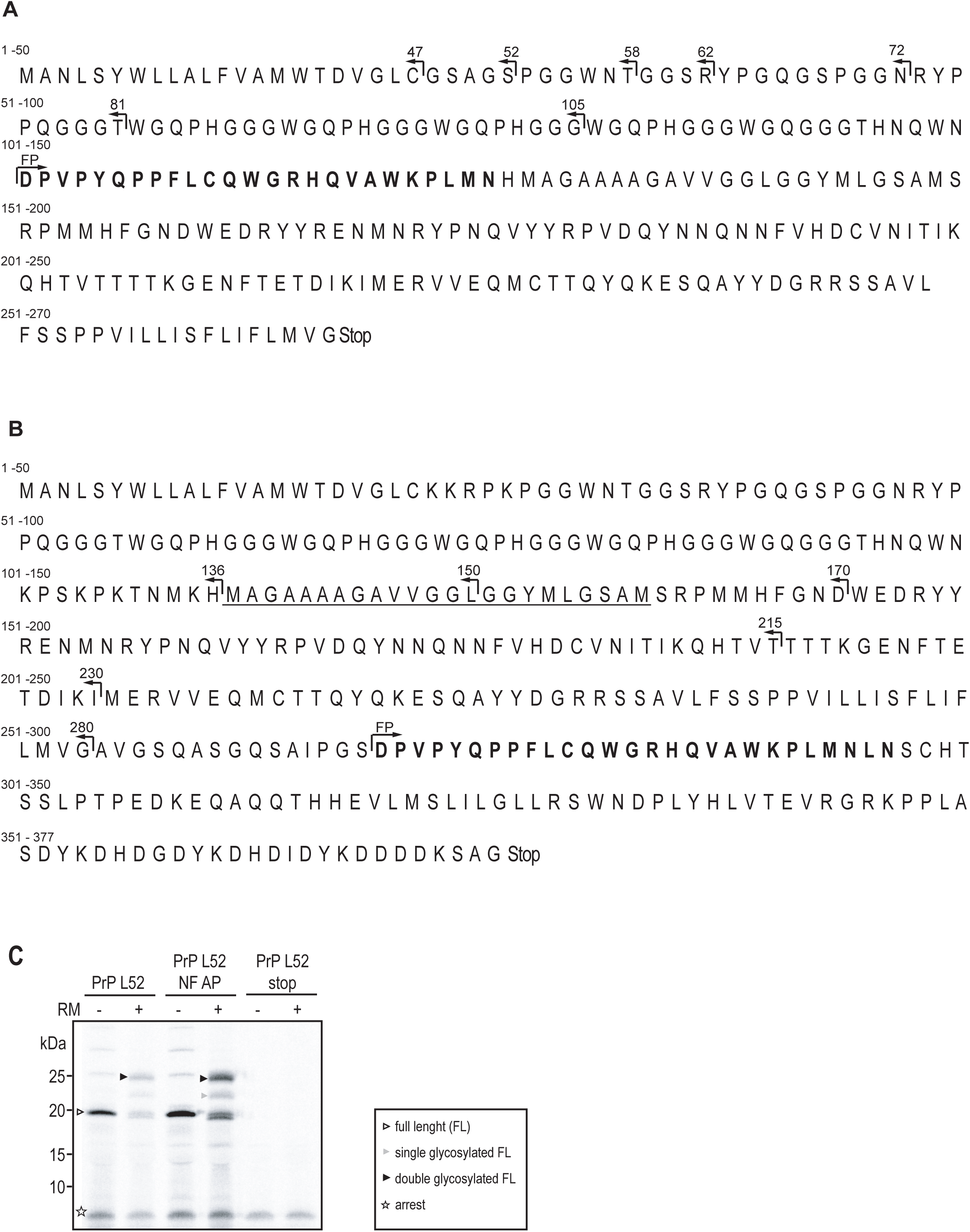
Amino acid sequences of basic constructs used in this study and a representative SDS--PAGE showing the migration pattern of a functional and a non--functional arrest peptide construct. The arrest peptide derived from mammalian X--box binding protein (Xbp1) is indicated in bold. The hydrophobic domain (HD) of PrP that can partition into the membrane is underlined. Small numbers in the front of the line indicate the position within the amino acids for each line. The positions for reverse primers binding to generate truncated variants of the two constructs are indicated by left facing arrows and the construct lengths generated until point of arrest. The binding site for the common forward primer is indicated by a right facing arrow and the abbreviation FP. (A) Primary sequence of PrP AP L126 construct used to generate all the shorter construct lengths as indicated by left facing arrows. (B) Primary sequence of PrP AP L295 construct used to generate all the shorter construct lengths as indicated by left facing arrows. (C) PrP constructs were *in--vitro* expressed in rabbit reticulocyte lysate in the presence of ^35^S--labelled methionine. Various populations were separated via SDS--PAGE and detected by autoradiography. The populations are marked as follows: double glycosylated full length (FL) protein: black filled arrow head; single glycosylated FL protein: grey arrow head; other FL fractions (including cleaved FL): empty arrow head; arrested protein fraction: empty asterisks. Identity of full lengths bands was confirmed by creating a weaker arrest peptide (AP), that had a lower stalling rate and led to more FL protein (PrP weaker AP). Identity of arrested bands was confirmed by replacing the last amino acid of the AP with a stop codon leading to protein product that migrates with the arrested proteins during SDS--PAGE (PrP stop).

**Supplementary figure 2.**
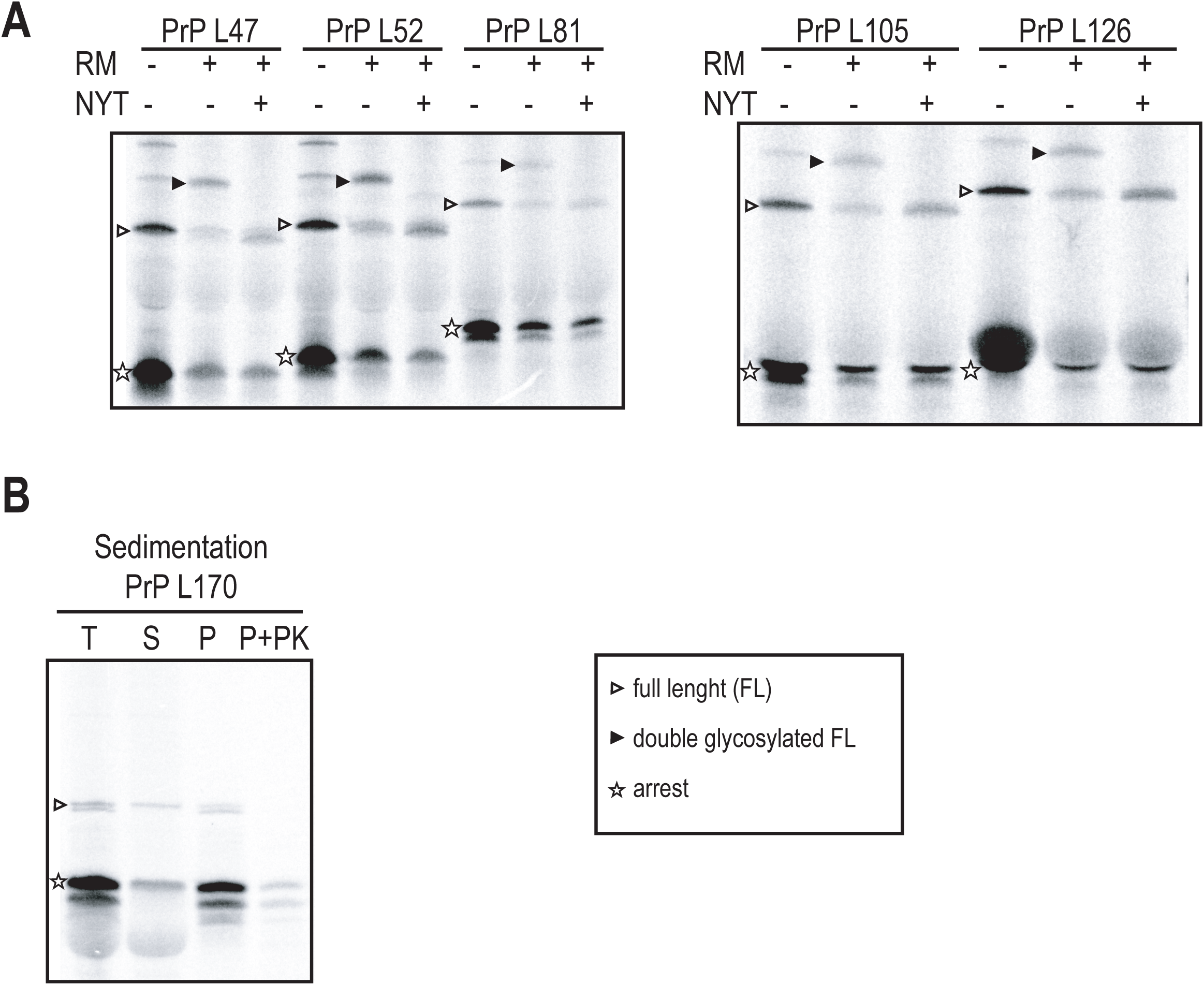
Pulled and arrested protein fractions can be discriminated by SDS--PAGE Various PrP constructs were *in--vitro* expressed in rabbit reticulocyte lysate in the presence of ^35^S--labeled methionine. Various populations were separated via SDS--PAGE and detected by autoradiography. The populations are marked as follows: double glycosylated full length (FL) protein: black filled arrow head; other FL fractions (including cleaved FL): empty arrow head; arrested protein fraction: empty asterisks; glycosylated arrested fraction: filled asterisks. (A) Identity of the glycosylated bands was confirmed by adding an inhibitor peptide (NYT) of N--linked glycosylation to the *in vitro* reaction. (B) Sedimentation on a sucrose cushion followed by proteinase K (PK) digestion of the pellet fraction reveals that PrP Ap L170 is an intermediate construct where cleavage of the arrest fraction (black arrow head) starts to become visible. The cleaved population is fully membrane associated and found in the pellet fraction (P), while parts of the uncleaved arrested protein can be found in the supernatant fraction (SN). Upon PK treatment both membrane--associated arrested populations appear to be equally protected, while the FL populations are not protected. The red arrow indicates the size where the Ntm--form of PrP can be found for longer construct lengths, but not for L170.

**Supplementary figure 3.**
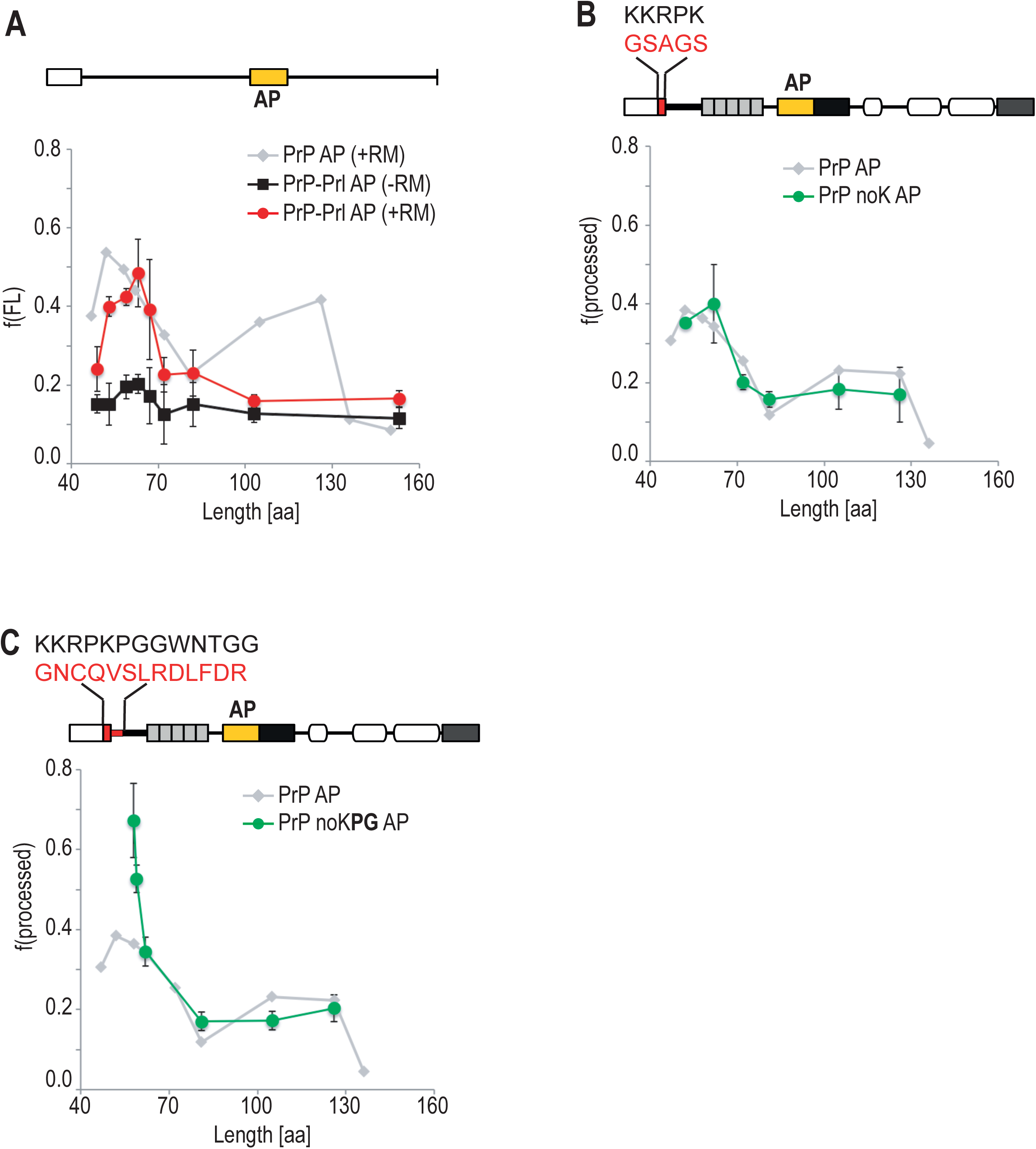
Force profiles for PrP--Prl and processed profiles for mutated variants of PrP. (A) Schematic representation of the constructs, top panel. Force profile for constructs that contains the PrP signal sequence in the background of prolactin (Prl) as a model protein with an arrest peptide introduced (PrP--Prl AP), bottom panel. Most data points are taken from our previous work and are shown again for illustration of the initial pulling event present for this construct (Kriegler *et al*., 2018). Data points show the average value of at least three independent experiments, error bars represent the standard deviation. (B) Schematic representation of the constructs, top panel. Fraction of processed full lengths protein (glycosylated and cleaved fractions) for the mutant PrP variant where the charged cluster is replaced by uncharged amino acids (PrP noK AP, green), bottom panel. The processed profile obtained for the wildtype PrP constructs is shown in grey. Data points show the average value of at least three independent experiments, error bars represent the standard deviation. (C) Fraction of processed full lengths protein (glycosylated + cleaved) for the mutant PrP variant were the charged cluster and 13 aa of the glycine and proline enriched region in PrP are replaced with a random sequence taken from Prl (PrP noK**PG** AP, green). The processed profile obtained for the wild type PrP constructs is shown in grey. Data points show the average value of at least three independent experiments, error bars represent the standard deviation.

**Supplementary figure 4.**
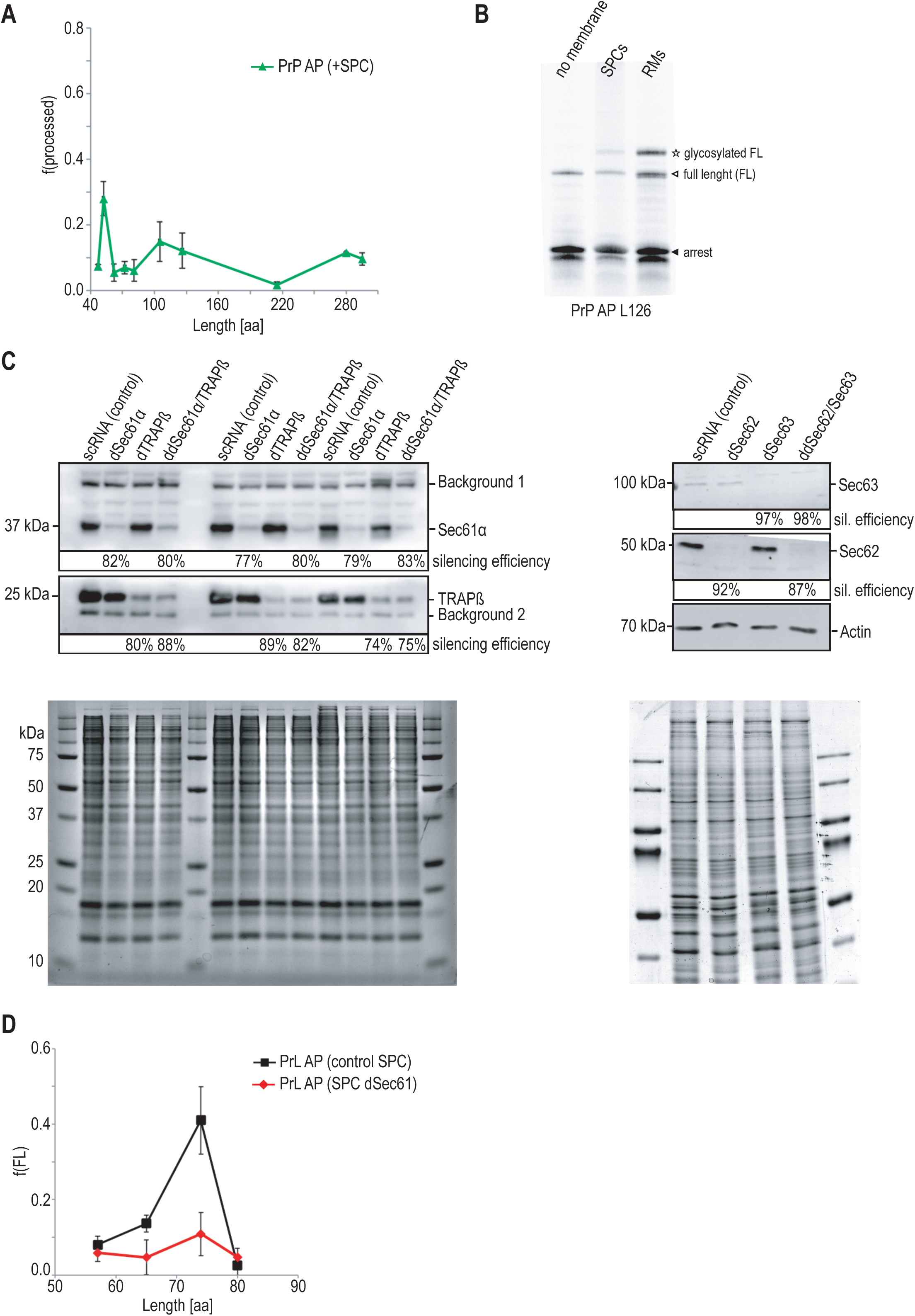
Semi--permeabilized cell system allows identification of components that have an effect on pulling events. (A) Fraction of processed protein (f(processed)) for PrP AP in presence of semi--permeabilized HeLa cells (SPCs) as ER membrane surrogate during *in vitro* protein expression. The graph represents the average of at least two individual datasets with the standard deviation shown as error bars; for constructs with large variability, more datapoints have been collected. (B) Representation of a representative experiment performed without membrane, or in presence of semi--permeabilized cells (SPCs) or dog pancreatic rough microsomes (RMs) for construct PrP L126 AP. (C) Example of western blots confirming the silencing efficiency for three experiments. The silencing efficiency was calculated from the band intensity in comparison to the noted background bands as loading control. Additionally, Coomassie stained gels also confirmed equal total protein amounts in SPC from silencing and control preparations. (D) Fraction of FL for Prl AP in the presence of SPCs treated with control siRNA (black) or SPCs treated with Sec61α siRNA (red). The graph represents the average of at least three individual datasets with the standard deviation shown as error bars.

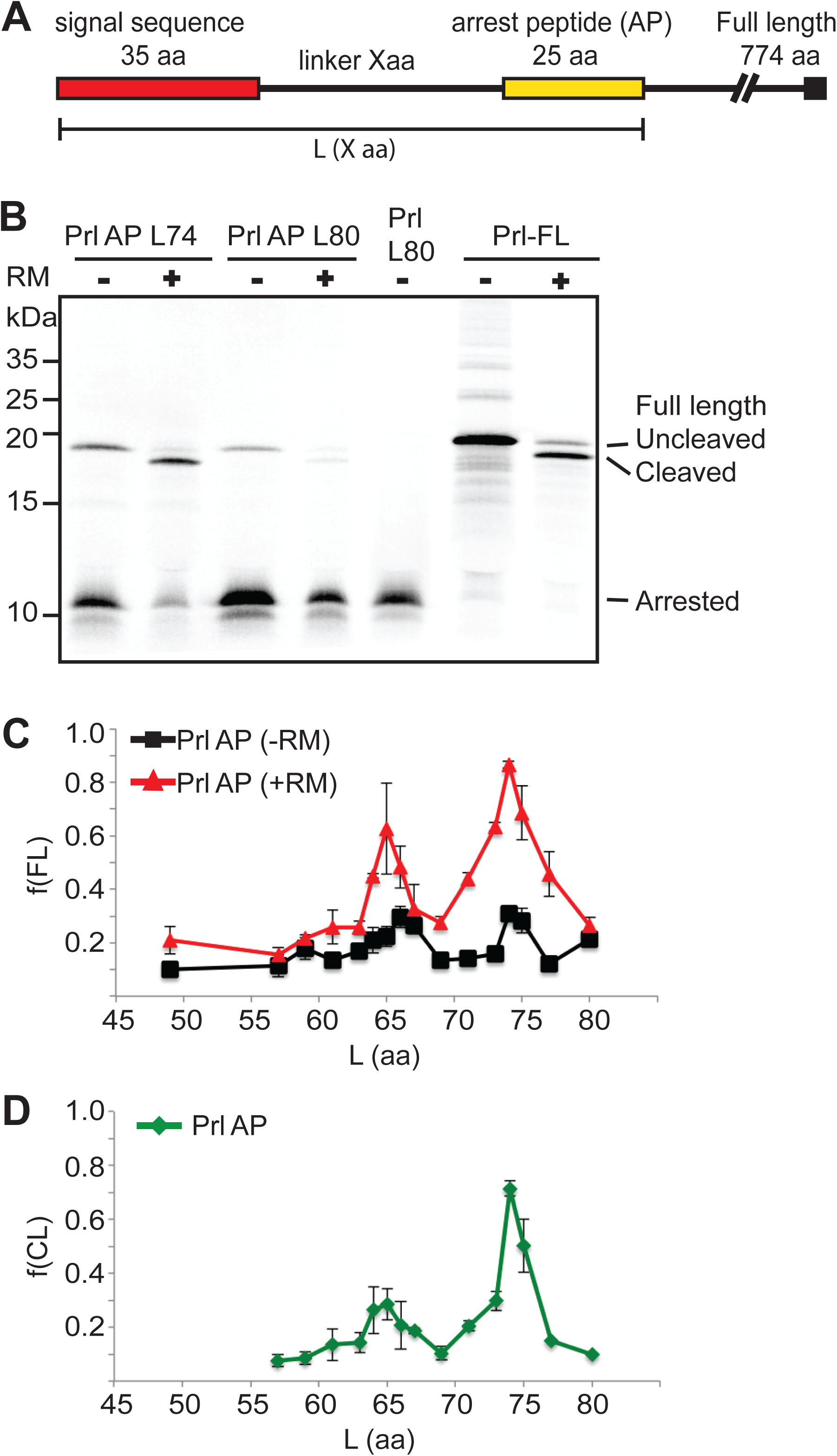

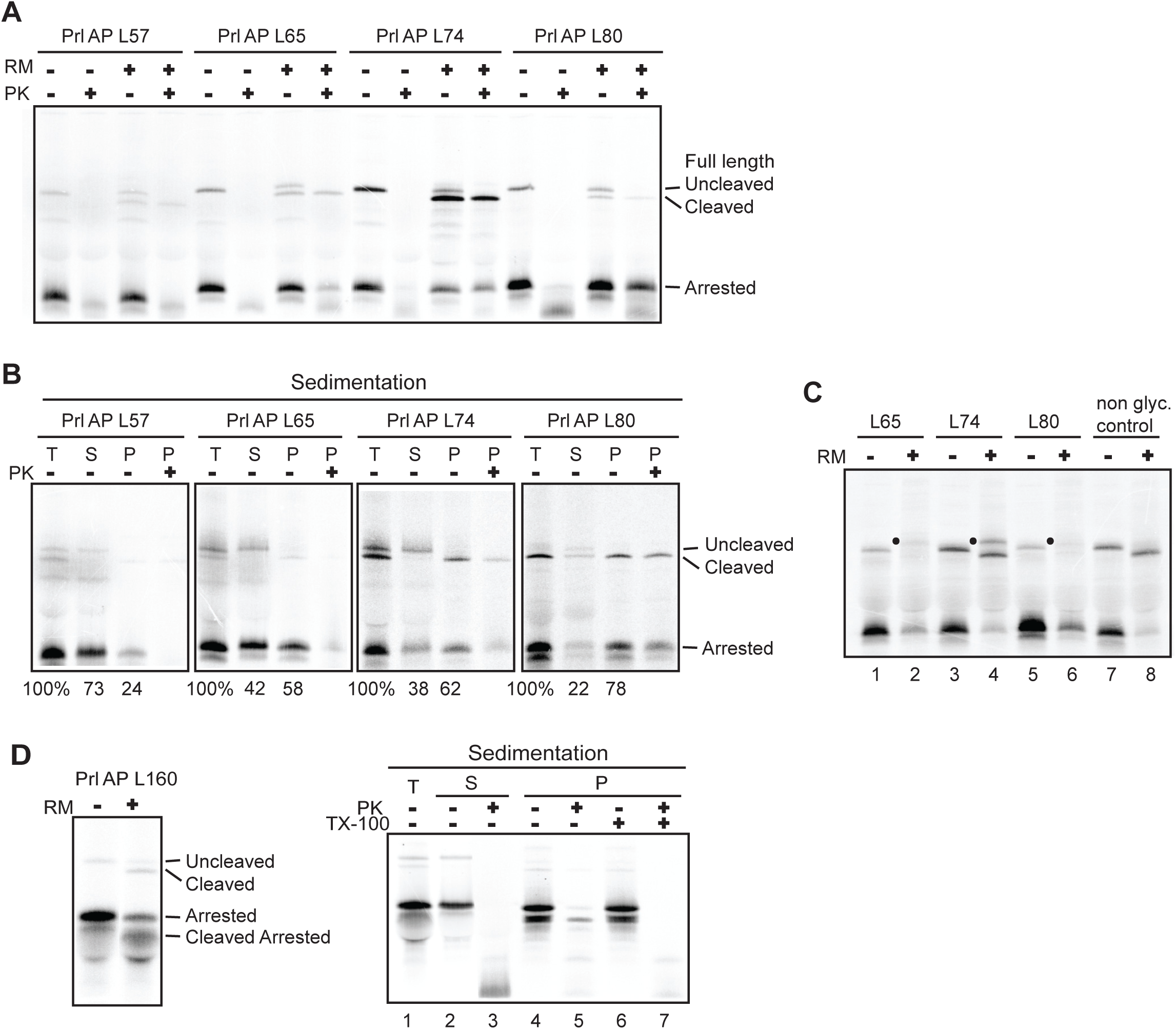

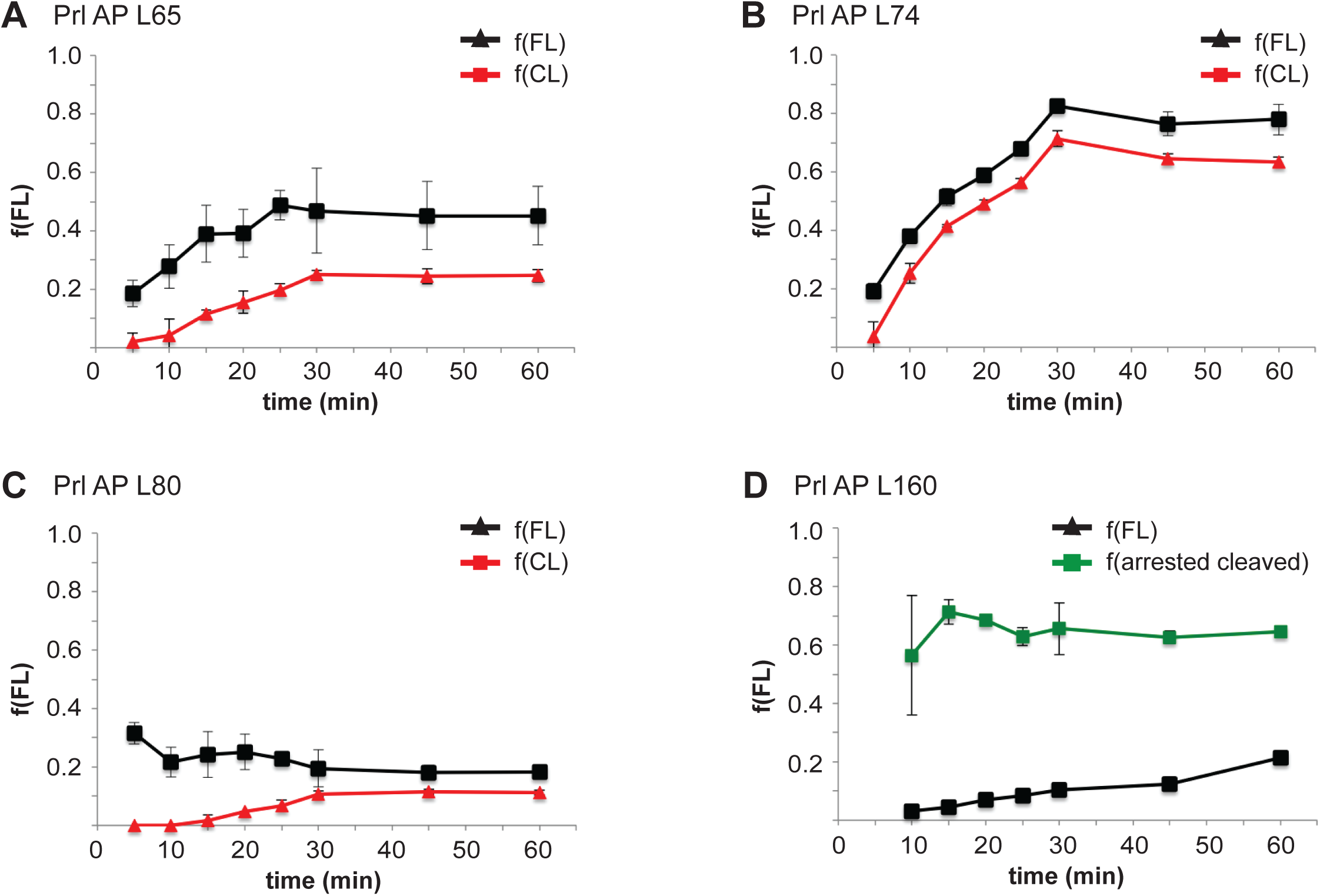

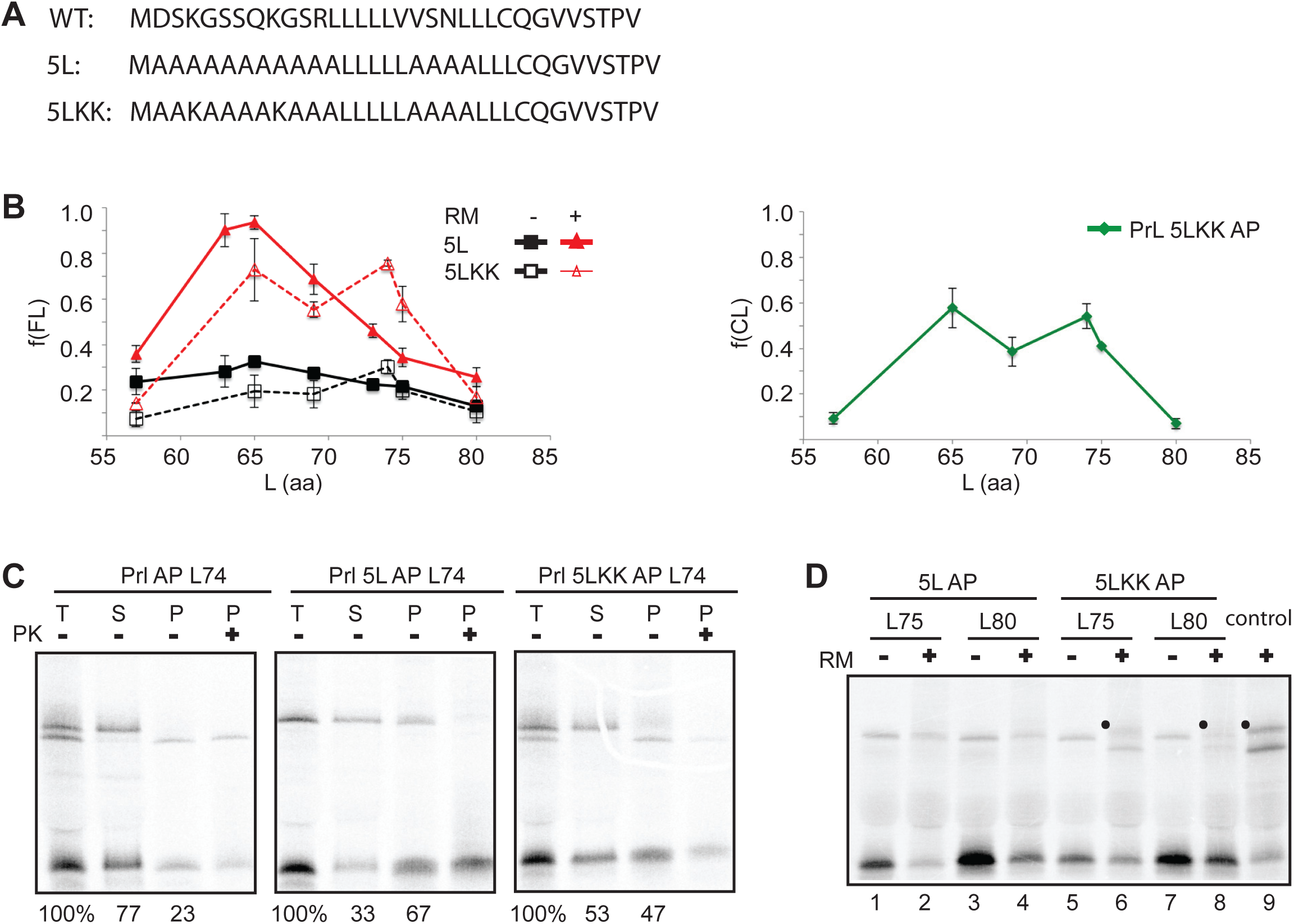

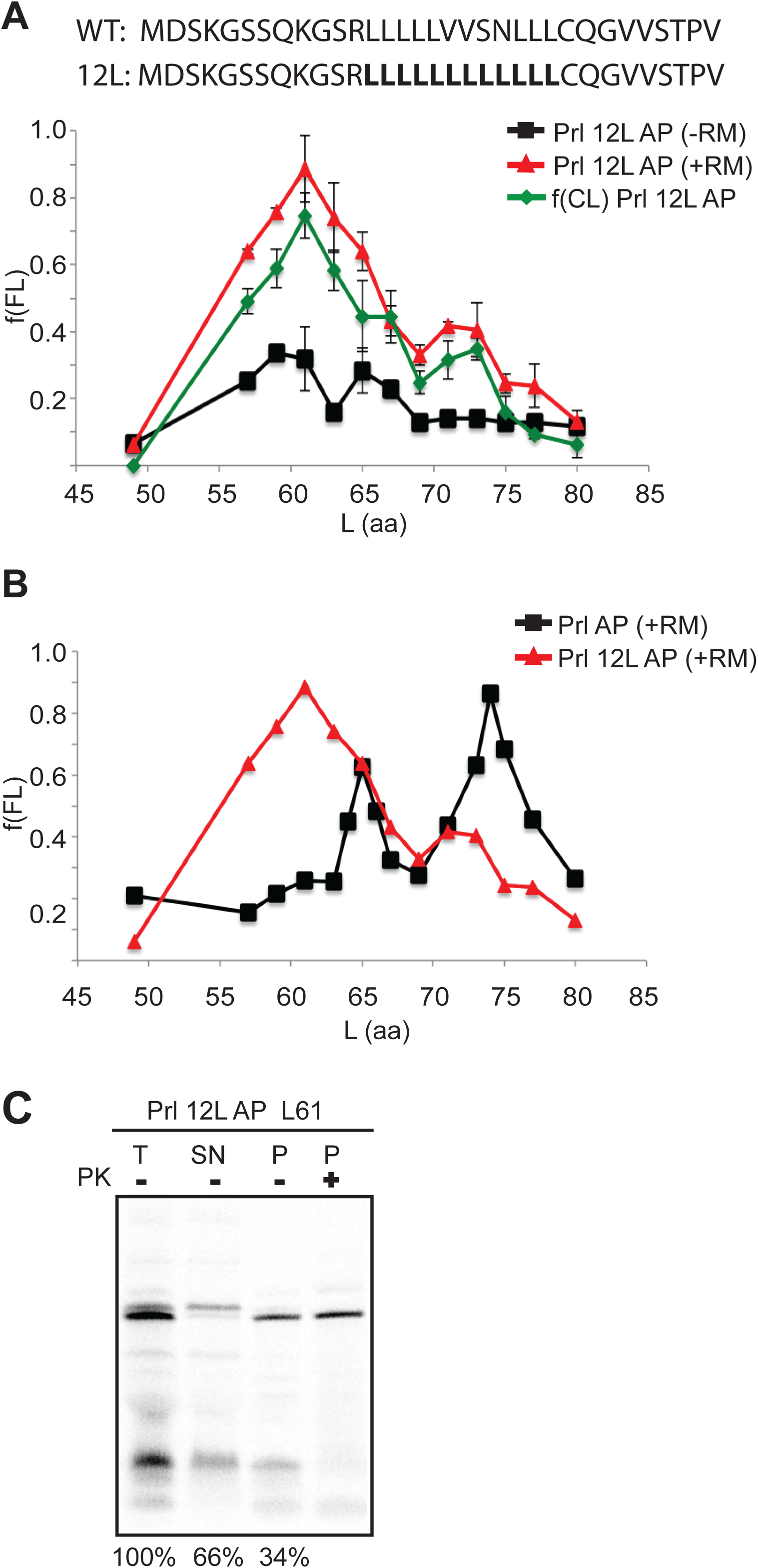

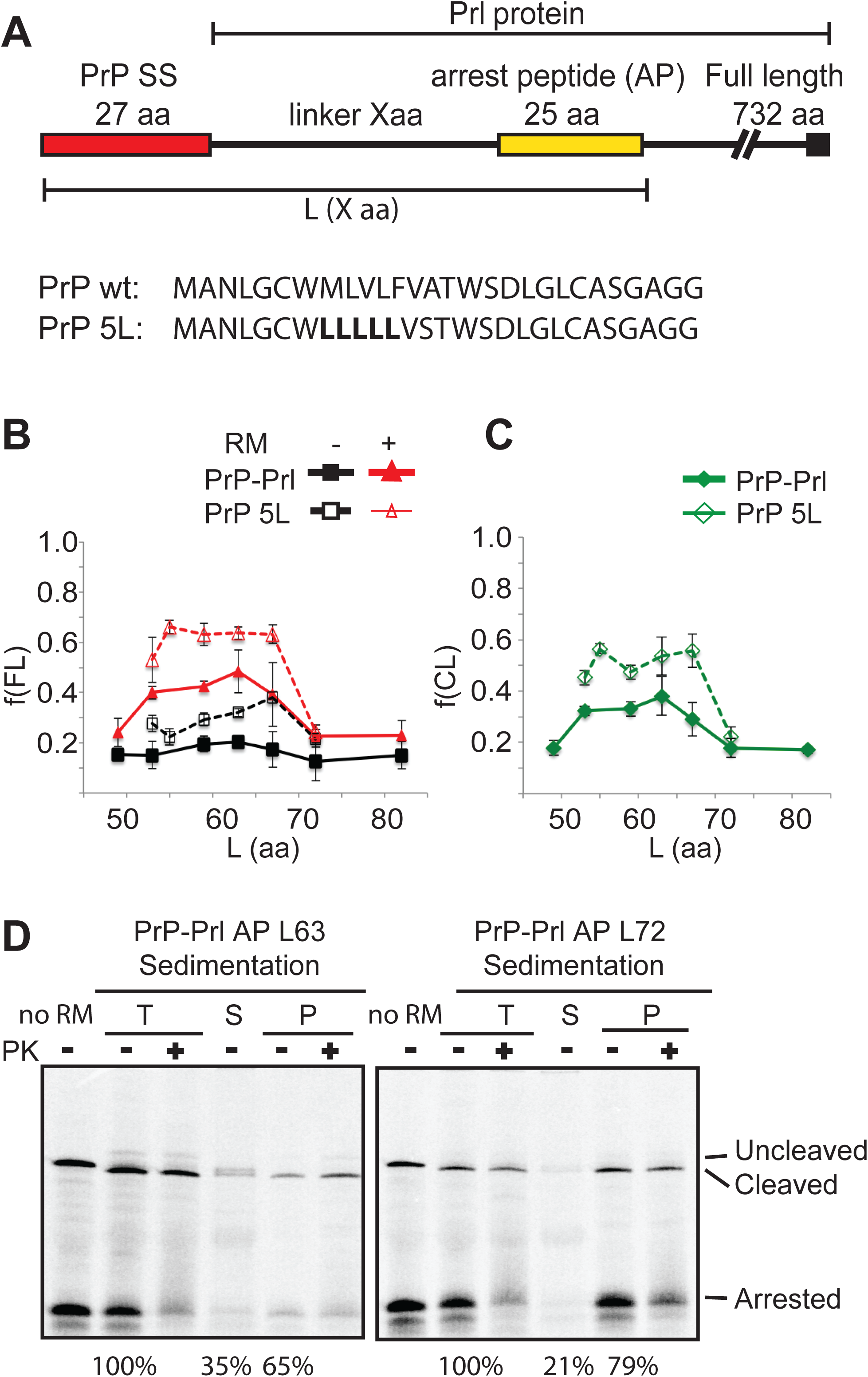

