## Supplementary figures and legends for "Prion Protein Folding Mechanism Revealed by Pulling Force Studies"

**Supplemantary Figure 1**

**A**

1 -50 (marked positions: 47, 52, 58, 62, 72)

M A N L S Y W L L A L F V A M W T D V G L C G S A G S P G G W N T G G S R Y P G Q G S P G G N R Y P

51 -100 (marked positions: 81, 105)

P Q G G G T W G Q P H G G G W G Q P H G G G W G Q P H G G G W G Q P H G G G W G Q G G G T H N Q W N

101 -150 (FP)

**D P V P Y Q P P F L C Q W G R H Q V A W K P L M N** H M A G A A A A G A V V G G L G G Y M L G S A M S

151 -200

R P M M H F G N D W E D R Y Y R E N M N R Y P N Q V Y Y R P V D Q Y N N Q N N F V H D C V N I T I K

201 -250

Q H T V T T T T K G E N F T E T D I K I M E R V V E Q M C T T Q Y Q K E S Q A Y Y D G R R S S A V L

251 -270

F S S P P V I L L I S F L I F L M V G Stop

**B**

1 -50

M A N L S Y W L L A L F V A M W T D V G L C K K R P K P G G W N T G G S R Y P G Q G S P G G N R Y P

51 -100

P Q G G G T W G Q P H G G G W G Q P H G G G W G Q P H G G G W G Q P H G G G W G Q G G G T H N Q W N

101 -150 (marked positions: 136, 150, 170)

K P S K P K T N M K H M A G A A A A G A V V G G L G G Y M L G S A M S R P M M H F G N D W E D R Y Y

151 -200 (marked position: 215)

R E N M N R Y P N Q V Y Y R P V D Q Y N N Q N N F V H D C V N I T I K Q H T V T T T T K G E N F T E

201 -250 (marked position: 230)

T D I K I M E R V V E Q M C T T Q Y Q K E S Q A Y Y D G R R S S A V L F S S P P V I L L I S F L I F

251 -300 (marked position: 280; FP)

L M V G A V G S Q A S G Q S A I P G S **D P V P Y Q P P F L C Q W G R H Q V A W K P L M N L N** S C H T

301 -350

S S L P T P E D K E Q A Q Q T H H E V L M S L I L G L L R S W N D P L Y H L V T E V R G R K P P L A

351 - 377

S D Y K D H D G D Y K D H D I D Y K D D D D K S A G Stop

**C**

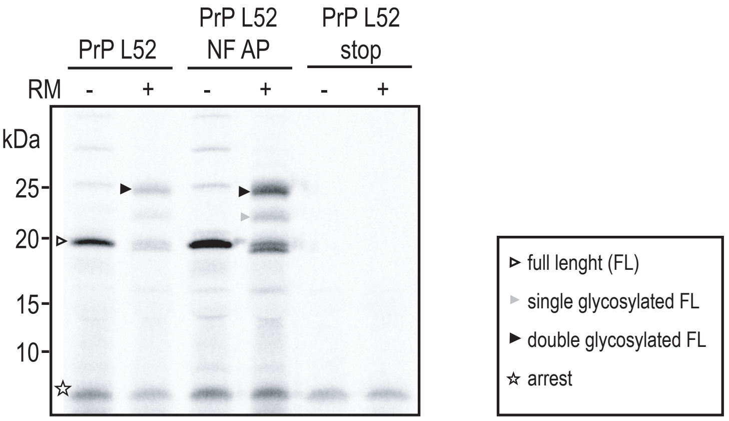

**Supplementary Figure 2**

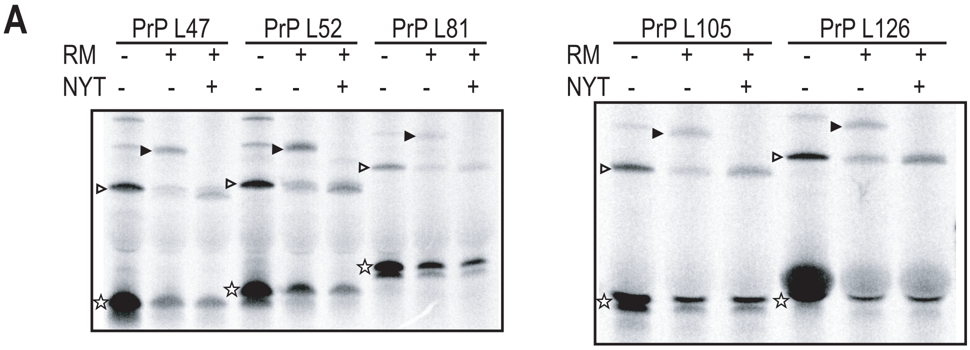


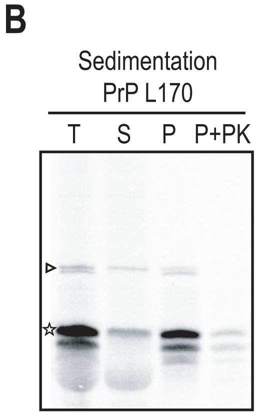


▹ full lenght (FL)

▸ double glycosylated FL

☆ arrest

**Supplementary Figure 3**

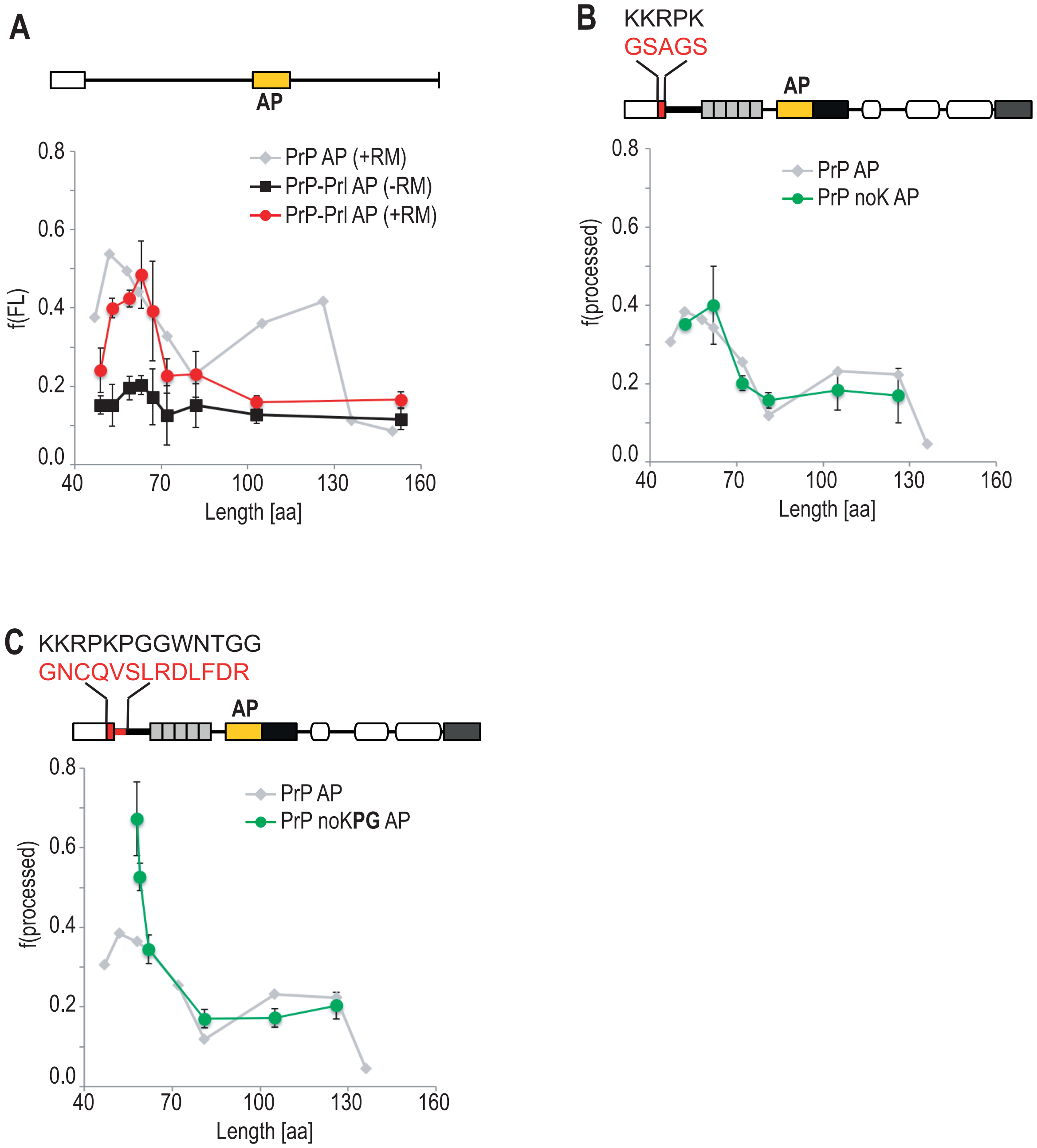

### Supplementary Figure 4

A

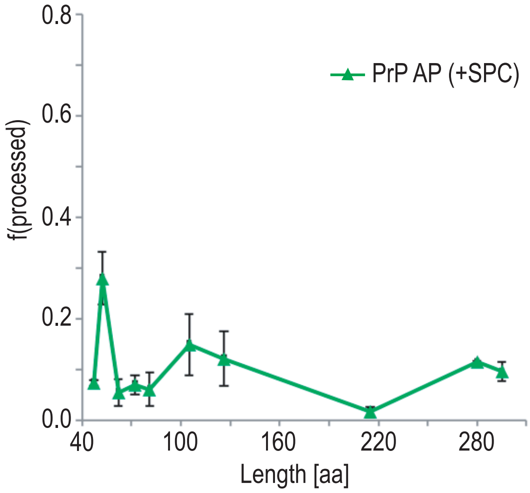


B

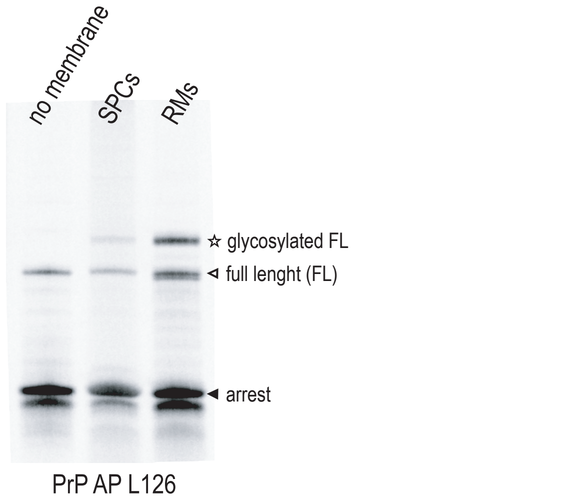


C

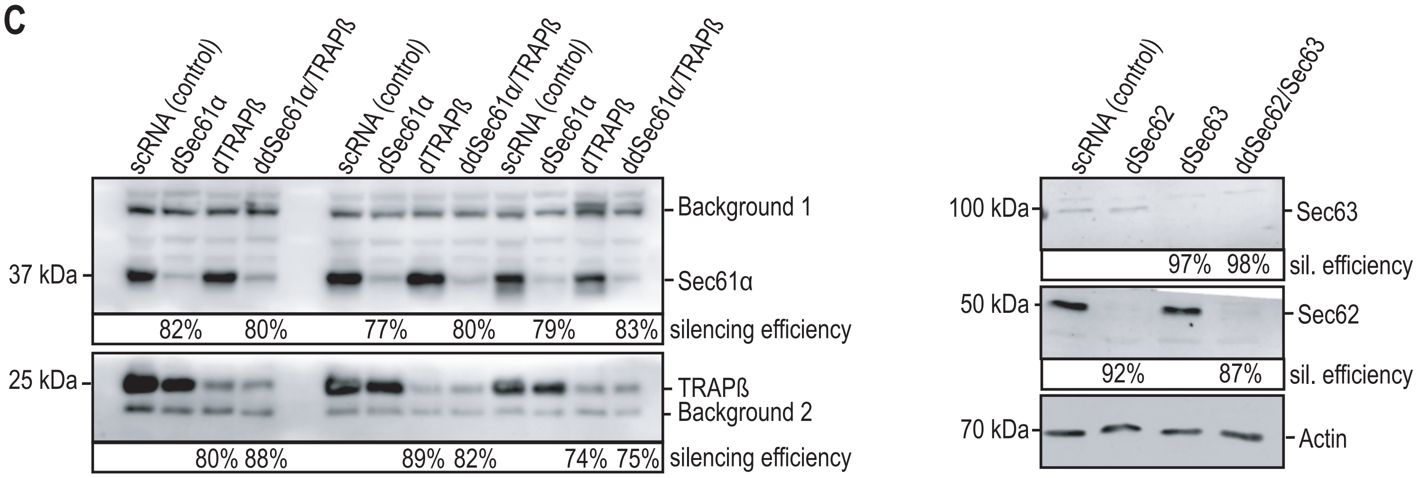


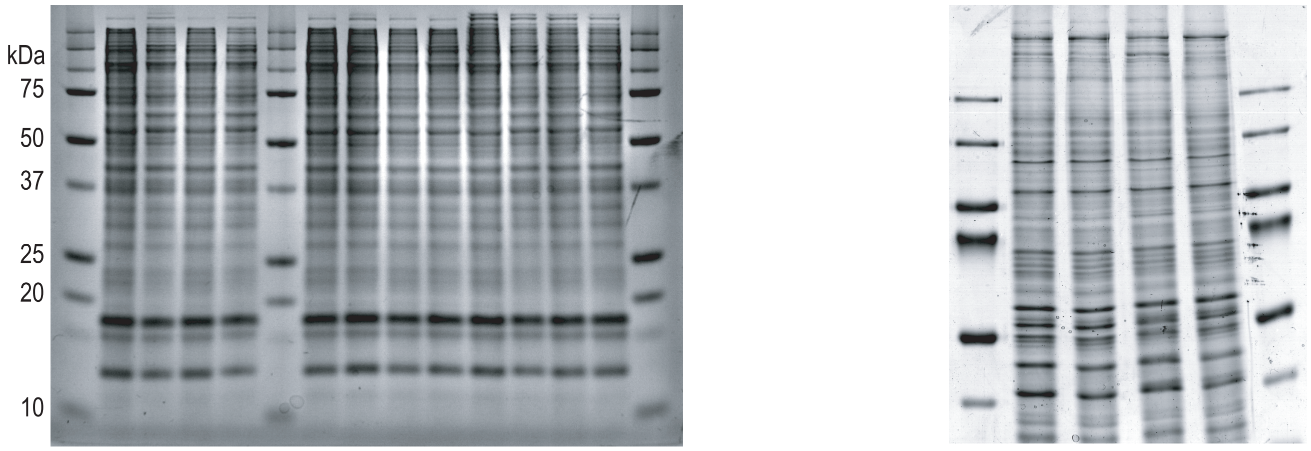


D

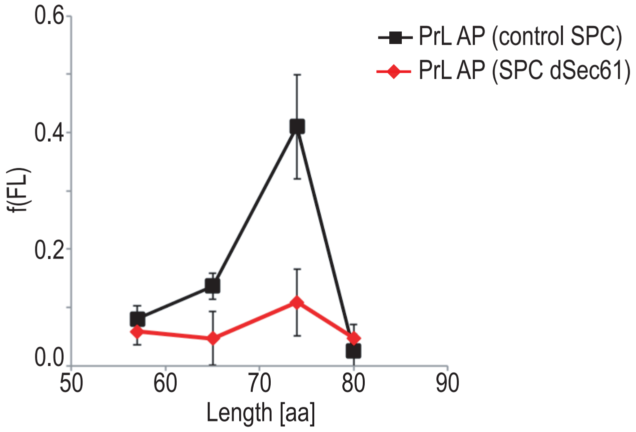
